# Walk-sum theoretical generators predict medial-temporal coupling driven by empirical cross-frequency generators: a novel joint cortical-subcortical approach in human sleep EEG

**DOI:** 10.64898/2026.04.18.719385

**Authors:** V. Kafetzopoulos, B. Kocsis

## Abstract

Large-scale brain activity can be described as walks on a connectome, and its dressed resolvent gives a compact walk-sum account of how signal flows among regions. A non-abelian extension of the resolvent introduces spectral-mode operators whose generators predict, empirically, that specific brain regions should display amplified cross-frequency phase-amplitude coupling with the rest of the brain. We refer to such regions as *empirical cross-frequency generators*. Methodologically, we introduce a joint cortical-subcortical source-imaging approach that combines a high-resolution cortical parcellation with a high-resolution subcortical parcellation in a single source-reconstructed M/EEG workflow, with validation gates at each step; to our knowledge this combination has not previously been documented as a validated pipeline. We compared cross-frequency phase-amplitude coupling during sleep in a 29-subject cohort against an awake rest-wake pool of 815 subjects. Eight subcortical parcels survived correction for elevated coupling in REM, all of them in the medial temporal lobe or the basal ganglia; the thalamus did not survive correction, although this null may reflect methodological sensitivity limits for deep structures rather than biological absence of thalamic coupling. We then turned the framework around and asked whether, given the known anatomical position of a target region — the nucleus reuniens, the obligate midline thalamic relay of the prefrontal-hippocampal circuit — the framework could predict its oscillatory function in advance. A densified 1 mm source grid over the Morel-defined reuniens mask, combined with a 68-parcel prefrontal target set, yielded a pattern compatible with the theta-and-beta directed PFC-Re-MTL routing suggested by the framework and by a prior rodent lesion study, with three distinct cells (N2 infraslow Re-to-temporal-pole, N1 beta Re-to-prefrontal, N1 beta Re-to-temporal) showing zero surviving pairs when extracted from a neighbouring thalamic parcel used as a negative control, arguing against beamformer point-spread as the sole explanation for the Re signature. A follow-up spectral-Granger analysis of cardiac inputs and ECG-derived respiratory surrogates showed cardiac leading the MTL hub across stages and bands, and the hub leading the slow respiratory envelope, in a direction that a purely mechanical amplitude-modulation estimate reproduces and that therefore may not be reducible to a heart-rate-variability artefact. A replication using direct intracranial recordings from the MNI Open iEEG Atlas (epilepsy patients, *= 1*−*10* per parcel) found cross-modal convergence: the left lateral amygdala ranked first in subcortical theta+beta coherence during N2, and generator-to-temporal coherence showed a stage-dependent band profile consistent with the scalp-level findings, although the intracranial results are limited by small and uneven coverage (see Limitations). A zero-hypothesis resting-state scan across three independent MEG cohorts (WAND *= 166*, COGITATE *= 100*, CamCAN *= 646*; total *= 912*) confirmed the left lateral amygdala as the top-ranked subcortical generator across all four datasets and three recording modalities, and demonstrated that the Re-temporal-pole coupling identified during sleep persists at rest. The walk-sum prediction is consistent with the data at two levels — both a data-driven hunt for generators and a pre-committed test of function-from-position — and the analysis pipeline documented here may offer an operational way to extend the framework-commitment recipe to any higher-order thalamic relay whose connectomic role is fixed.

## 1. Introduction

The network view of the brain rests on two complementary analytical traditions: structural connectomics, which treats the brain as a fixed wiring diagram, and functional connectivity, which measures statistical dependencies between regional time series [25-27]. The walk-sum resolvent framework bridges them by treating the connectome as a discrete walk space on which signals propagate, and summing over all walks to obtain a dressed resolvent that captures both direct and indirect paths between any pair of nodes. In its scalar form the resolvent assigns one complex number per pair per frequency, and a truncated sum over walks of moderate length already reproduces empirical resting-state functional connectivity to within scan-rescan ceiling.

A natural and algebraically clean extension promotes the scalar propagator at each edge to an operator that acts on a multi-band spectral state. The propagators no longer commute in general, and the resulting dressed resolvent takes the form of a path-ordered product: a non-abelian analogue of the scalar theory. The generators of this extended theory are abstract algebraic objects, the generators of the spectral-mode Lie algebra, and we refer to them throughout as *theoretical generators* to distinguish them from the empirical brain regions they predict. Two empirical consequences follow. First, the natural three-point function of the non-abelian theory is the cross-bispectrum [11-16, 22], whose antisymmetric part is provably zero for instantaneous volume-conduction mixing and is therefore a leakage-immune directional measure of coupling [16]. Second, the existence of loci where this three-point function is amplified, loci we call *empirical cross-frequency generators*, is a structural prediction of the theoretical generators: such regions should exist, their signature should be most visible where competing cortico-cortical activity is reduced, and the appropriate observable for finding them is a cross-frequency phase-amplitude coupling measure such as Gaussian-copula mutual information [9] or bicoherence [16]. The theory does not predict which anatomical regions these empirical generators are. That is an empirical question, and it is the question this paper addresses.

Sleep provides a convenient low-arousal probe. It strips away a large portion of the awake cortical ensemble while leaving the subcortical machinery that supports theta, delta and spindle oscillations broadly intact [31, 32, 35, 39, 40]. If a region is a coupling generator then its phase-amplitude coupling with the rest of the brain should grow in sleep relative to awake rest, rather than shrink as overall cortical power drops. The experimental test is concrete: compute phase-amplitude coupling per subcortical parcel in sleep, compare with matched awake rest, and identify the parcels that survive a fair multiple-comparisons correction. We emphasise that sleep EEG is chosen here as a probe rather than as the scientific object of interest. The walk-sum prediction is about coupling generators, not about sleep; any sufficiently noise-reduced low-arousal state should in principle work, and sleep is one convenient such state that is easy to acquire and easy to stage. What we seek to test is the theory’s prediction; what the specific sleep-state findings may offer in addition is a window on how the generators behave in a biologically meaningful state.

The second contribution of this paper is the analysis toolset itself. Running the test end-to-end requires (i) full-brain source reconstruction with both cortical and subcortical coverage from scalp EEG, (ii) a parcellation dense enough to give per-region statistics but not finer than the effective spatial resolution of the source reconstruction, (iii) a non-parametric phase-amplitude coupling estimator that tolerates non-Gaussian signals, and (iv) a null-validation procedure that guards against over-interpretation of the per-parcel survivor count. Individual components of this workflow are well established [1-10, 16], but to our knowledge no published pipeline combines a high-resolution cortical parcellation with a high-resolution subcortical parcellation in a single source-reconstructed M/EEG analysis with validation gates at each step. We document that pipeline here alongside the findings, and we expect it to be useful in applications beyond the specific scientific question posed in this paper.

## 2. Theoretical framework

Let be an × connectivity matrix with spectral radius ()<*1* after normalisation, and let denote angular frequency. The walk-sum resolvent is

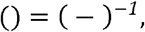

which admits the Neumann expansion

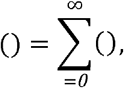

with each term summing all walks of length on the graph induced by, phase-delayed by. The non-abelian extension promotes the scalar propagator at each edge to a spectral-mode operator acting on a vector in an -band state space. The dressed resolvent becomes a path-ordered product over hops,

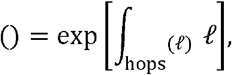

and the underlying algebra is () rather than (*1*). Equation (3) is a generating object from which empirical predictions follow; we do not evaluate it directly. Its empirical consequence is that the natural three-point function of the theory is the cross-bispectrum

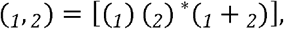

which measures how the phase of a slow mode at site couples to the amplitude of a fast mode at site through a third site. The magnitude normalisation, bicoherence, is

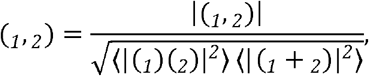

and the leakage-immune antisymmetric variant of Chella et al. [16] is

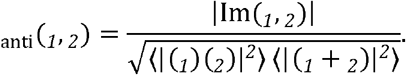

The spectral-mode operator is the object that implements hierarchical cross-frequency routing in the extended theory: because slow phases gate fast amplitudes in the non-abelian algebra, the operator that mixes spectral modes at an edge is precisely the operator that would carry any hierarchical top-down control of that edge. Cross-frequency phase-amplitude coupling is therefore the natural observable for the integrity of, and within-band power and the topological connectivity are observables for the two logically independent aspects of the framework (local dynamics and scaffold connectivity); a perturbation confined to would reduce cross-frequency coupling while sparing power and, and the three readouts can in principle be measured independently.

For practical phase-amplitude coupling estimation we use the Gaussian-copula mutual information of Ince et al. [9]. Each marginal is rank-transformed through the inverse Gaussian CDF,

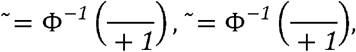

and the mutual information is then evaluated in closed form on the Gaussian copula,

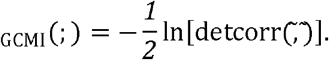

GCMI is asymptotically equivalent to Shannon mutual information for jointly Gaussian variables and is a lower bound for general non-Gaussian joints. We use it as our primary phase-amplitude coupling metric; bicoherence results are reported in the supplementary material as a reviewer reserve.

For directionality of the identified couplings we use spectral Granger causality [18, 19, 21]. Writing for the spectral density matrix and Γ for the associated transfer function,

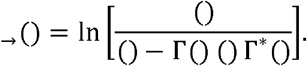

The factorisation is performed non-parametrically in the implementation we use [18]. See [20] for the related information-theoretic transfer-entropy formulation.

## 3. Methods

### 3.1 Datasets

The sleep cohort was the AnPhy-Sleep dataset [49], 29 healthy adults recorded at the Montreal Neurological Institute with high-density scalp EEG (83 electrodes, ∼71 usable EEG channels after reference and ground removal) at 250 Hz across a full night of polysomnography, with expert-scored hypnograms. The awake rest-wake comparison pool combined two independent cohorts processed through an identical source-reconstruction pipeline: the Welsh Advanced Neuroimaging Database (WAND) [50] (169 subjects, resting-state MEG from Cardiff University Brain Research Imaging Centre) and the Cambridge Centre for Ageing and Neuroscience (Cam-CAN) cohort [47, 51] (646 subjects, resting-state MEG), for a total of = *815* awake subjects. A third independent MEG cohort, the COGITATE dataset [103] (= *100* subjects, resting-state MEG), was used in the zero-hypothesis resting-state replication of 4.8 alongside WAND and CamCAN (total = *912* for that analysis). Exclusions were applied on pre-specified quality criteria alone: the number of qualifying sleep epochs per stage, the number of bad channels after kurtosis thresholding, and the presence of required physiological channels. No subject was excluded on the basis of the analyses below.

### 3.2 Preprocessing

EEG data were notch-filtered at 50 Hz and bandpassed to 0.5-80 Hz. Bad channels were detected by variance and kurtosis thresholding and interpolated. Channel names carrying a reference suffix (for example Fp1-Ref or O2-M1) were sanitised to their base forms so that the standard 10-20 electrode template would match. Without this step only a small fraction of the 71 channels is recognised and the source reconstruction silently collapses to a near-trivial leadfield; we flag this as a non-obvious but critical preprocessing step for AnPhy-format data.

### 3.3 Source reconstruction and whole-brain parcellation

Source reconstruction used a linearly-constrained minimum-variance beamformer [1] implemented in MNE-Python [2]. The head model was the fsaverage boundary-element model with the Destrieux-labelled cortical surface [4] as the anatomical reference. The parcellation was the union of the 800-region Schaefer 17-network cortical atlas [5, 6] and the 54-region Tian S4 subcortical atlas [7] (comprising 10 hippocampal, 16 thalamic, 4 amygdalar, 8 putamen, 8 caudate, 4 nucleus accumbens and 4 globus pallidus parcels), giving 854 parcels in total across cortex and subcortex.

We use the 800-parcel cortical variant because its mean parcel volume (∼0.6 cm^*3*^) sits at the spatial resolution floor of source-reconstructed 71-channel scalp EEG and co-locates at that grain with the Tian S4 subcortical parcels; a sensitivity analysis across alternative cortical and subcortical parcellations is in Supplementary §S7.

Applying the full beamformer projection to 854 parcels on a continuous 10 min recording at 250 Hz materialises an intermediate tensor of several hundred gigabytes on our host, well beyond available host memory. We therefore chunked the beamformer application in 30 000-sample blocks (2 min) with no overlap and concatenated the reconstructed sources. Verification at the parcel level: all 854 parcels had non-empty vertex lists on the fsaverage surface; hemisphere balance was preserved after chunking; no parcel received zero time points; and subcortical leadfields for the hippocampal and amygdala parcels passed the magnetoencephalographic deep-source sanity criterion of Pu et al. [48] of non-zero sensitivity above simulation noise floor. That review provides evidence that non-invasive magnetic sensors can recover hippocampal rhythms; scalp EEG is expected to be harder than MEG for deep structures, and we therefore read the medial-temporal and basal-ganglia parcel time series as spatially approximate rather than anatomically precise, with caveats flagged in the Discussion.

### 3.4 Sleep staging and epoching

Stage N1, N2 and REM epochs were extracted from the expert-scored hypnograms. For parity with the awake comparison cohort, whose typical usable recording length is approximately 10 min, we capped each sleep stage at 20 non-overlapping 30-second epochs. Subjects with fewer than 10 qualifying N1 epochs, or fewer than 20 qualifying N2 or REM epochs, were excluded from the corresponding stage analysis. After these exclusions: _*1*_ = *22*, _*2*_ = *29, = 29*.

### 3.5 Cross-frequency coupling

For each parcel we computed Gaussian-copula mutual information [9] between the phase of a lower-frequency band and the amplitude of a higher-frequency band, across four band pairs (theta-gamma, alpha-gamma, theta-beta, alpha-beta). Slow-band phase and fast-band amplitude were extracted by forward-backward FIR bandpass filtering followed by the Hilbert analytic signal. Before the GCMI estimator we downsampled each subject’s joint to 4000 samples to satisfy the approximate IID assumption of the underlying k-nearest-neighbour density estimator [10] without discarding within-subject information. The effective degrees of freedom of the joint are set by the slower band, and the subject-level GCMI mean is stable to within 5% under a ±*20*% variation in the downsample count. Per-parcel neighbour-mean GCMI was defined as the mean GCMI over all pairs touching that parcel.

### 3.6 Generator identification

Per subject we computed the log ratio log(AnPhy_stage_/pool_awake_) at each parcel, and tested against zero across subjects by a sign-flip permutation test (10 000 permutations, seed 0). Benjamini-Hochberg false discovery rate correction [24] was applied within each (stage, CFC-pair) family of 54 subcortical parcels. A subject-level bootstrap 95% confidence interval (10 000 resamples, seed 0) accompanies each point estimate for effect-size context; the interval excludes unity wherever a parcel survives FDR correction.

### 3.7 Spectral Granger causality

Spectral Granger causality was computed with the non-parametric state-space solver [18] as used in the MVGC framework [19]. Three bands were used with band-adapted epoch lengths: infraslow (0.05-0.5 Hz, 60 s epochs with 50% overlap), theta (4-8 Hz, 20 s non-overlapping) and beta (13-30 Hz, 20 s non-overlapping). The maximum Granger lag was 20 samples (80 ms at 250 Hz). The generator seed set was the eight subcortical parcels identified in the per-parcel test of §3.6; the cortical reference set was the 16 left anterior temporal parcels that survived FDR correction when the eight-generator network was used as a seed in a separate network-level null against random 8-parcel subcortical selections. The autonomic channel set was strictly cardiac (ECG1, ECG2) and the ECG-derived respiratory surrogates described in §3.8; muscle and ocular channels were excluded as non-autonomic by construction. Per-pair paired t-tests of (forward GC − reverse GC) across subjects were corrected with BH-FDR within each (stage, band, direction-pair) family.

### 3.8 ECG-derived respiration

The sleep dataset does not contain a respiratory belt. To obtain a respiratory-band observable we derived three surrogate signals from each ECG lead. EDR-A, R-peak amplitude modulation from thoracic-axis rotation during breathing, is a purely mechanical consequence of the chest moving during inspiration and expiration and does not depend on the R-R intervals at all. RSA, the respiratory-band component of heart-rate variability, is by definition heart-rate variability in the respiratory band and therefore captures any cardiovagal respiratory modulation. EDR-BW, baseline-wander EDR, is obtained by bandpassing the raw ECG directly to the respiratory range and therefore shares low-frequency content with the raw ECG waveform [45, 46]. R-peaks were detected with a median-absolute-deviation threshold on a 4-Hz high-passed ECG and a 0.4-second refractory interval. All three surrogates were bandpassed to 0.05-0.5 Hz with a 4th-order Butterworth SOS filter. EDR-A and RSA provide complementary non-HRV and HRV estimates; EDR-BW is reported with caution and treated as a confound rather than a clean mechanical control (Supplementary §S5).

### 3.9 Replication pipeline and validation gates

Figure 1 gives the per-subject pipeline as a seven-step flow with a validation gate printed alongside each step. The pipeline is intended to be reproducible from Figure 1 and the descriptions above by a reader familiar with MNE-Python and the two parcellations; the scripts used for the present analysis are available from the corresponding author on request.

**Figure 1.**
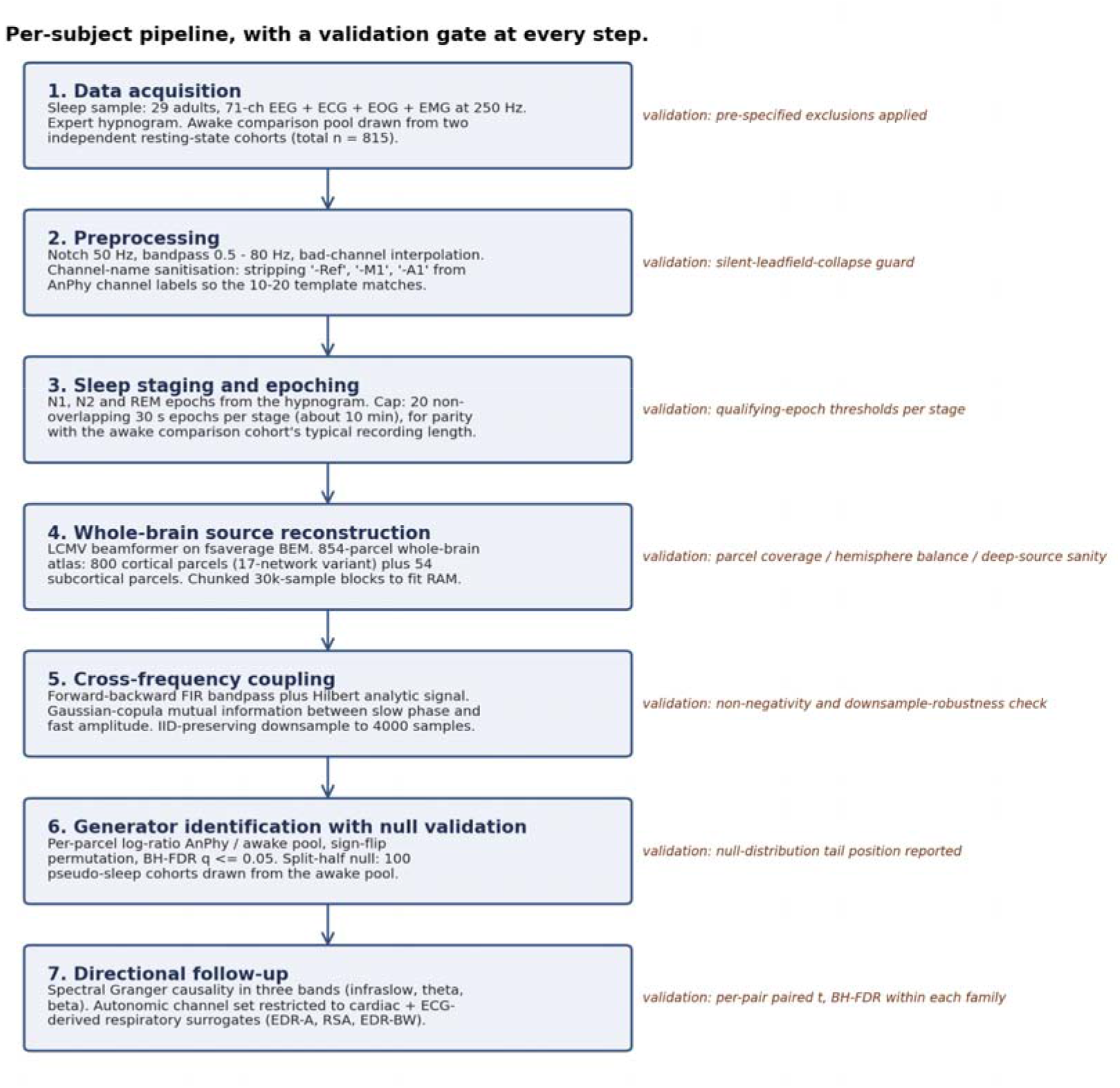
Per-subject analysis pipeline in seven steps, with a validation gate at each step. The critical non-obvious steps are Step 2 (channel-name sanitisation, which otherwise silently destroys the leadfield), Step 4 (chunked beamformer projection to the 854 cortical-plus-subcortical parcels, with the deep-source sensitivity sanity check of Pu et al. [48]), and Step 6 (split-half null-distribution validation of the FDR-surviving parcel count, without which the per-parcel survivor count is easy to over-interpret).

## 4. Results

### 4.1 Eight subcortical parcels, all non-thalamic

Per-parcel elevation of theta-beta GCMI in REM, tested against the awake rest-wake pool, identified eight subcortical parcels surviving BH-FDR correction at (Table S1, Figure 2, Figure 3). The surviving set was three hippocampal subdivisions (HIP-head-l-lh, HIP-head-m2-lh, HIP-body-lh), three amygdalar subdivisions (lAMY-lh, lAMY-rh, mAMY-lh), one right anterior globus pallidus (aGP-rh) and one left putamen/ventral pallidum (PUT-VP-lh). Effect sizes ranged from a ratio of 1.43 (PUT-VP-lh) to 2.43 (HIP-head-l-lh), and every per-parcel 95% subject-level bootstrap confidence interval excluded unity. Zero thalamic parcels survived in any stage or CFC pair, and no subcortical parcel survived in N1 or N2 for any CFC pair. However, the absence of thalamic survivors should be interpreted with caution: source reconstruction from 71-channel scalp EEG has limited sensitivity to deep midline structures, and the thalamic null may reflect a methodological sensitivity floor rather than a genuine biological absence of thalamic coupling (see Limitations). The REM theta-beta cell carries the bulk of the generator signal in this dataset.

**Figure 2.**
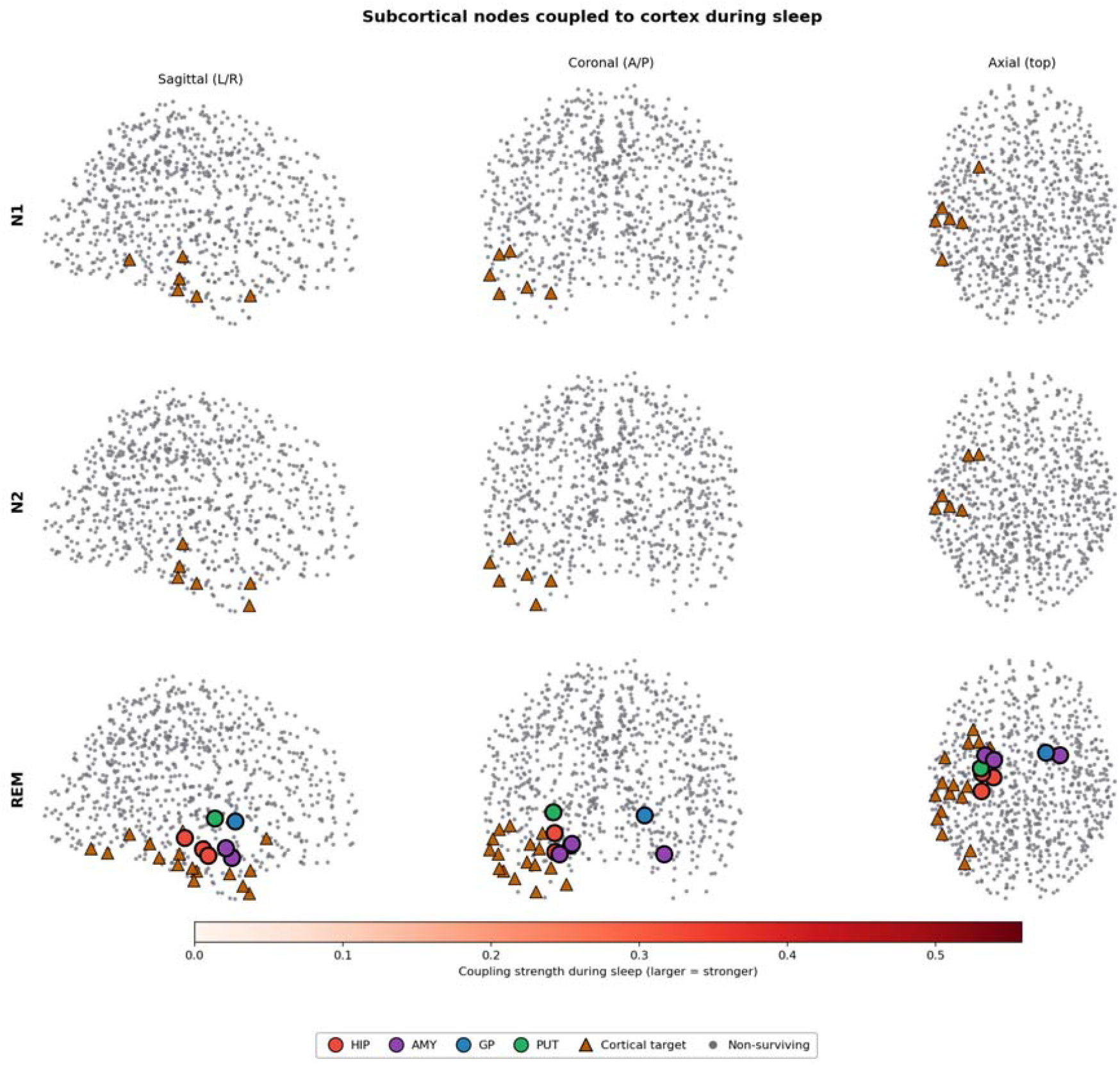
Subcortical coupling generators across sleep stages. Each row is a sleep stage (N1, N2, REM) and each column is an orthogonal glass-brain projection (sagittal, coronal, axial). Coloured dots mark the centroid of each subcortical parcel whose theta-beta GCMI survived BH-FDR correction against the awake rest-wake pool. Dot colour indicates the anatomical group (HIP, AMY, GP, PUT). In N1 and N2 no parcel survives. In REM, eight parcels survive: three hippocampal, three amygdalar, one globus pallidus and one putamen / ventral pallidum. No thalamic parcel survives in any stage, although this may reflect the limited spatial resolution of scalp EEG for deep midline structures (16 thalamic parcels of 54 subcortical). Background: fsaverage cortical surface, transparent.

**Figure 3.**
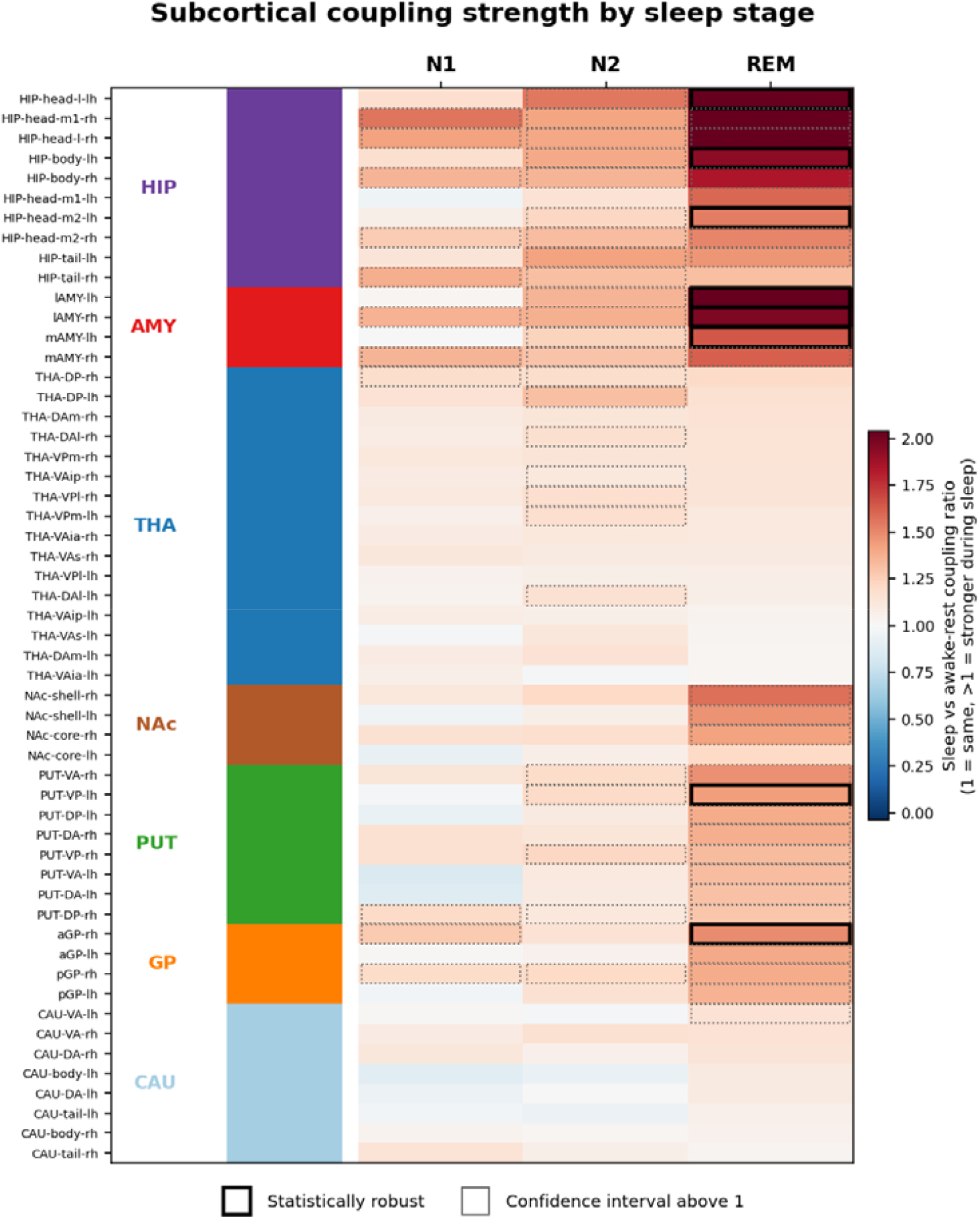
Stage-by-parcel heatmap of the log(sleep / awake-rest) theta-beta GCMI ratio. Rows are the 54 subcortical parcels grouped by anatomical class (HIP, AMY, GP, PUT, CAU, NAc, THA) and hemisphere; columns are the three sleep stages. Cell shade encodes effect size. Dots mark cells surviving per-parcel BH-FDR correction within the (stage, CFC-pair) family. Thalamic rows are uniformly near the null; MTL and BG rows are elevated in REM; both lAMY parcels on the left hemisphere survive in REM.

A split-half null validation, constructed by drawing 100 pseudo-sleep cohorts of size 29 from the awake pool and running the identical analysis pipeline on each, placed the observed REM theta-beta survivor count at the 98.0th percentile of the empirical null distribution (Table S2). This is robust, but not extreme, and we refrain from numerical claims about the exact number of surviving parcels; the stable qualitative finding is that the set is non-empty, non-thalamic, and anatomically concentrated in the medial temporal lobe and the basal ganglia.

### 4.2 Cortical targets in the left anterior temporal lobe

Taking the eight generators as a fixed network and asking which cortical parcels show elevated theta-beta GCMI to that network against a random 8-parcel subcortical null (10 000 iterations), we found 6 cortical targets surviving FDR correction in N1, 6 in N2 and 16 in REM. Five cortical parcels were common to all three stages: two default-mode-B temporal parcels (Temp_2, Temp_4), one more inferior default-mode-B parcel (Temp_10), and two limbic-A temporal-pole parcels (TempPole_10, TempPole_11). Network-level enrichment was strongest for limbic-A temporal pole in REM (6 of 27 parcels, 22%), which overlaps the left anterior temporal hub of the semantic system [29], and for default-mode-B temporal cortex (4 of 52 parcels, 8%), the ventral temporal component of the default-mode network [28]. All surviving targets were in the left hemisphere, consistent with the left-lateralised seed set.

### 4.3 Cardiac input drives the hub in every sleep stage

Spectral Granger causality, with the autonomic channel set restricted to cardiac channels, identified a cardiac hub direction (throughout §4.3-4.6 “the hub” denotes the eight-parcel generator set identified in §4.1, i.e., three hippocampal, three amygdalar, one globus-pallidus and one putamen/ventral-pallidum parcel) surviving per-pair BH-FDR correction in every combination of sleep stage (N1, N2, REM) and band (infraslow, theta, beta) we tested. Best per-pair values ranged from to, best per-pair values were below, and between 10 and 14 of the 16 cardiac-to-generator pairs survived correction in each cell of the stage-by-band table. ECG2 was the dominant driver; ECG1 was consistent but weaker. Figure 4 summarises the directionality at the stage-pooled level.

**Figure 4.**
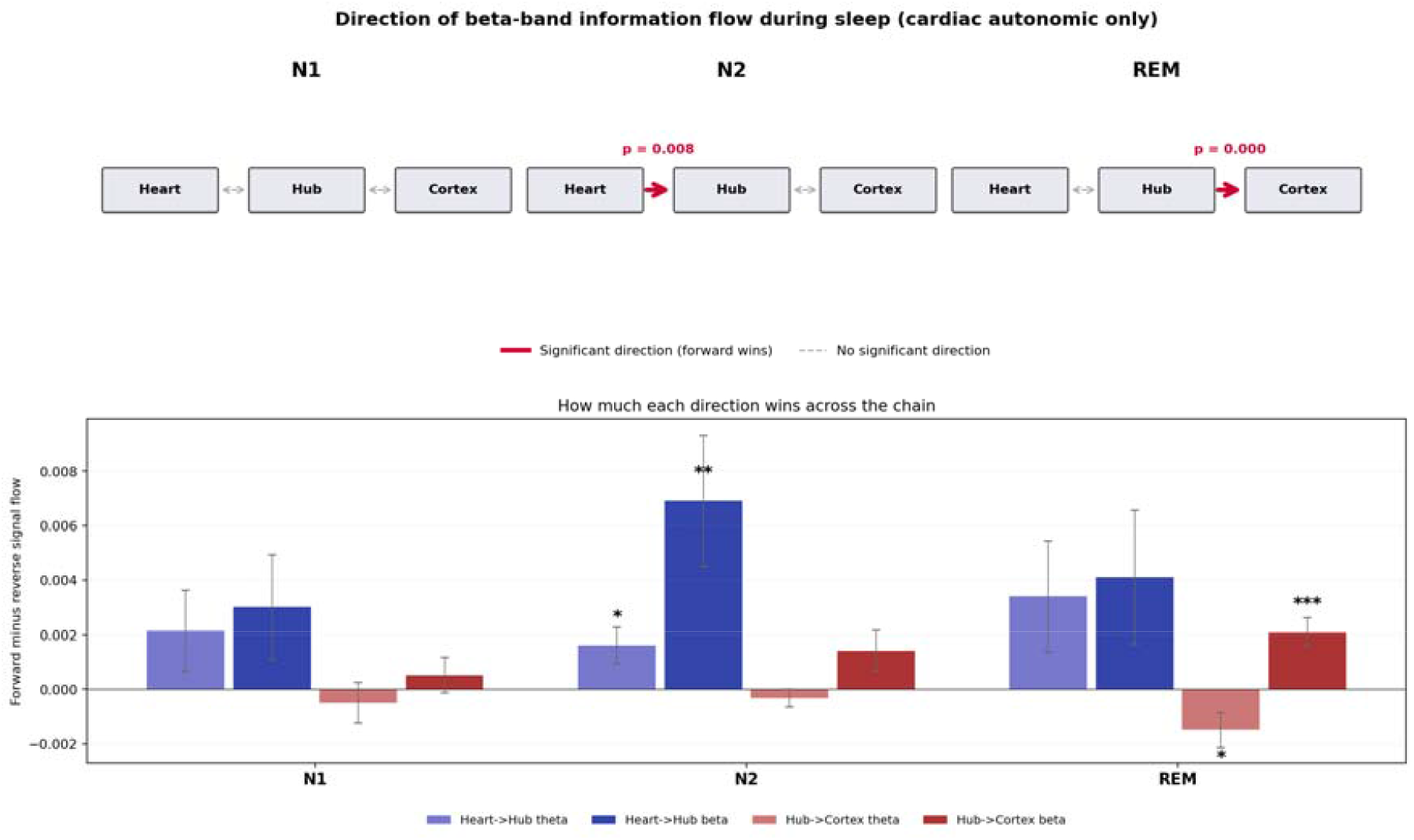
Directional information flow in sleep. Top: schematic, one column per sleep stage. Bold red arrows mark directions significant at the stage-pooled level after per-pair averaging; dashed grey arrows mark non-significant directions. Bottom: grouped bar chart of the stage-pooled forward-minus-reverse Granger value per direction pair and band (theta and beta shown; infraslow is reported in the text). Bars are mean across subjects; error bars are standard errors; stars denote uncorrected paired-t significance (,,). The cardiac-to-hub direction is significant at the stage-pooled level in N2, and the hub-to-cortex direction is significant in REM beta.

**Figure 5.**
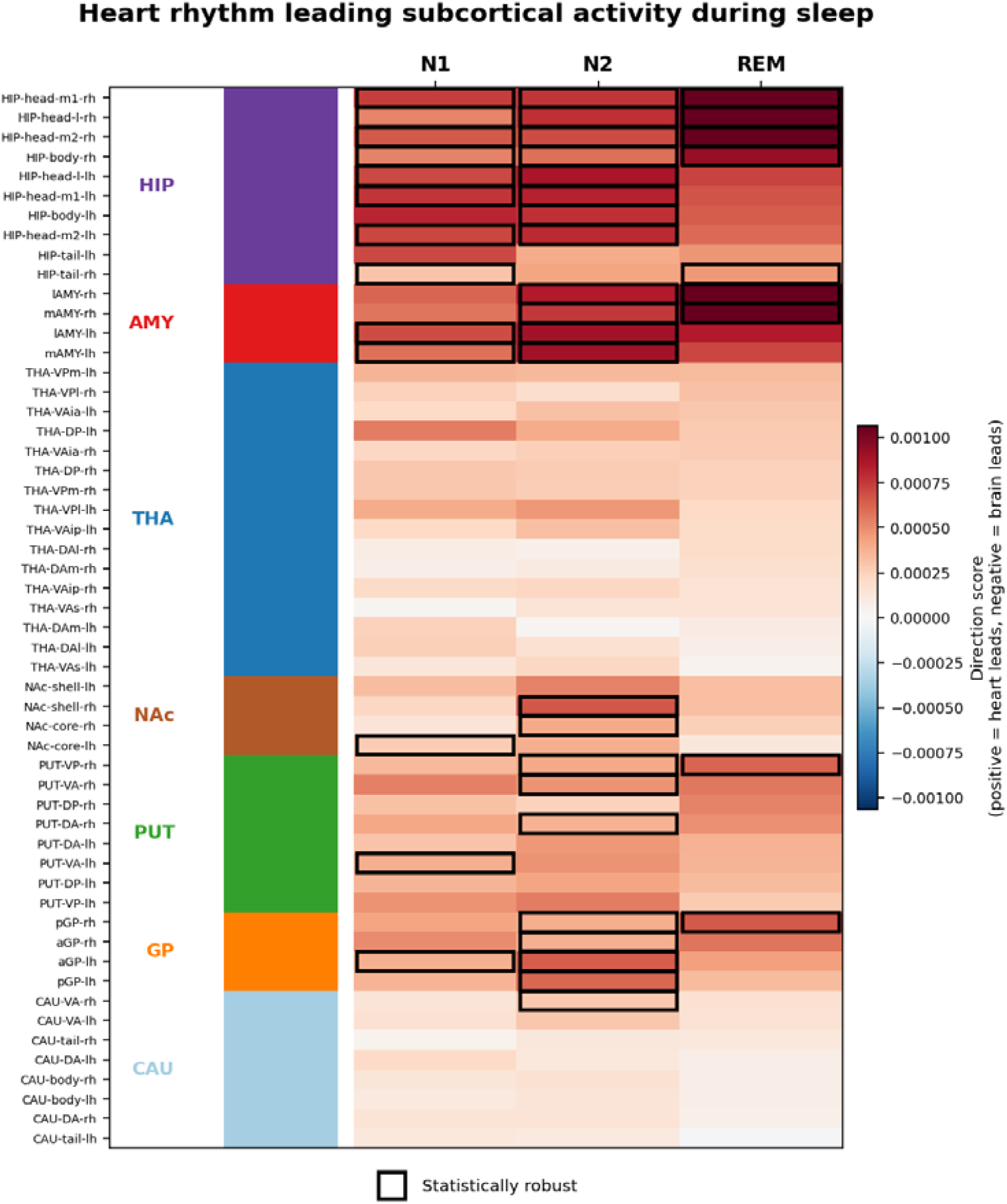
Per-parcel directional score of cardiac subcortical coupling across sleep stages. Rows are the 54 subcortical parcels grouped by anatomical class; columns are the three sleep stages. Cell shade is the subject-mean of direction score; positive (red) means cardiac phase leads subcortical amplitude, negative (blue) the reverse. Dots mark cells surviving per-parcel BH-FDR correction within the (stage, slot) family. The medial-temporal and basal-ganglia rows are robustly positive across all three stages; thalamic rows are uniformly near the null.

### 4.4 Hub-to-cortex directionality: REM beta and N2 infraslow

In REM beta, the forward direction (hub cortex) produced 24 pairs surviving per-pair BH-FDR correction. The top forward pairs were hippocampal-body, medial-amygdala and ventral-pallidum seeds projecting to the left anterior insula, the left default-mode-B temporal cortex, and the left temporal pole. N1 beta showed a weaker three-pair hippocampus left anterior insula signature. N2 infraslow (0.05-0.5 Hz) produced the single largest per-pair effect in the analysis: 20 forward-direction pairs surviving correction, with the top pair being HIP-head-m2-lh limbic-A temporal pole (forward minus reverse difference), and a general pattern of MTL seeds projecting to the left temporal pole.

### 4.5 Framework-committed prediction and test of the nucleus reuniens

The analyses of §4.1-4.2 identify MTL and basal-ganglia cross-frequency generators and their cortical targets without any anatomical prior, and the directional analyses of §4.3-4.4 characterise the hub’s autonomic inputs and cortical outputs. We now turn the framework around and ask the complementary question: given an anatomical target whose *position* in the mammalian connectome and whose *known inputs and outputs* are already fixed, can the framework predict the oscillatory function it should display, and are the data consistent with that prediction? The target is the nucleus reuniens (Re). Re sits at the ventral midline of the thalamus between prefrontal cortex and the ventral hippocampus, receives a dense projection from medial prefrontal cortex, and is the obligate relay through which prefrontal cortex reaches CA1, because CA1 projects directly to PFC but PFC has no direct CA1 projection [52, 55-59, 63]. Re is therefore the single edge carrying the top-down half of the prefrontal-hippocampal dialogue [60-62, 64, 65]. Under the commitment of §2, an edge whose anatomical role is to carry hierarchical top-down routing is precisely an edge whose spectral-mode operator should show non-trivial cross-frequency structure; and under the rat-lesion result of Kafetzopoulos et al. [52] we know the bands at which the corresponding PFC-HPC coherence is Re-dependent — theta, beta and low gamma — and the bands at which it is Re-independent — delta and high gamma. Putting the two together, the framework commits, in advance of any human data, to three predictions: (i) Re should show directed coupling with PFC on the input side and with MTL / anterior temporal cortex on the output side; (ii) the coupling should live at theta and beta; (iii) the effect should be state-specific and strongest in the consolidation-relevant stages (N2 and REM), rather than in N1. Our aim in this section is *not* to discover a new anatomical relay — Re is already fully characterised by rodent tract-tracing — but to show that the framework, combined only with the known anatomy and the joint cortical-subcortical source-imaging approach of §3, generates a concrete and testable prediction about oscillatory function. This is an illustration of how the framework may be used going forward to interrogate any region whose position and inputs/outputs are known.

#### Clean Re isolation via the Morel histological thalamic atlas

The 54-region Tian subcortical atlas used in §4.1-4.4 does not contain a reuniens-specific label, and its closest ventral-midline thalamic parcel (THA-VAia) overlaps the Morel Re mask [53, 54] at only ∼*1*% of voxels per hemisphere. We therefore constructed an augmented mixed source space that inserts a densified 1 mm-resolution volumetric grid inside the reuniens mask of the Morel histological atlas of the thalamus (Krauth et al. [53]; medioventral nucleus, corresponding to rodent Re): 16 source points in the left hemisphere and 13 in the right. A second LCMV beamformer pass projected the 71-channel EEG through this augmented head model and produced a Re time series, averaged per hemisphere, that samples only tissue inside the Morel Re mask. Cortical targets were simultaneously expanded to include 68 prefrontal parcels from the same 800-region Schaefer atlas (26 orbitofrontal, 19 medial prefrontal, 10 ventromedial prefrontal, 3 frontopolar, 7 salience medial prefrontal, 2 mid-cingulate and 1 anterior cingulate), so that the canonical PFC-Re-HPC circuit of [52] could be tested in its directed human form. The Tian THA-VAia bilateral pair was carried through the same pipeline as a negative control: VAia’s centroid sits about 5.3 mm lateral and dorsal to Morel Re, so its beamformer point-spread at the ∼*5* mm subcortical grid overlaps Re substantially. Any signal genuinely living in Re should partially leak into VAia; any signal exclusively present in the Re extraction but absent in VAia cannot be explained by this leakage. Full details of the negative-control analysis are in Supplementary §S8.

#### Results

Spectral Granger causality on the Morel Re time series yielded a pattern compatible with the directed PFC-Re-MTL routing suggested by [52] at the committed bands (Figure 6, panel A). In stage N2, Morel Re showed directed coupling with the expanded prefrontal target set at both theta (13 FDR-surviving pairs) and beta (23 FDR-surviving pairs), within their (stage, band, direction-pair) family. A corresponding N2 infraslow Re ⟶ prefrontal cell (20 FDR-surviving pairs) was also detected but is shared with the THA-VAia negative-control extraction at near-identical effect sizes (Supplementary §S8) and is therefore not claimed here as a Re-specific signature, although it is consistent with a broader ventral-midline-thalamic-to-prefrontal routing to which Re contributes. The largest forward-minus-reverse differences at theta were into medial prefrontal and dorsal-control prefrontal parcels (Morel_RE-lh ⟶ RH_ContB_PFCmp_1-rh, RH_DefaultA_PFCm_10-rh, RH_ContB_PFCmp_2-rh, all ≥*3*.*5*, ≤ *0*.*031*). The largest forward-minus-reverse differences at beta were into orbital and medial prefrontal parcels (Morel_RE-lh ⟶ LH_LimbicB_OFC_3-lh, RH_DefaultA_PFCm_1-rh, both ≥*4*.*3*, ≤ *0*.*022*). These are the theta and beta prefrontal signatures predicted by the commitment and by the Re lesion pattern of [52]; their absolute specificity to Re versus the broader ventral-midline-thalamic compartment is partial at the source resolution available here, with Re dominating the VAia negative control in surviving pair count (13 vs 6 at theta; 23 vs 13 at beta) but not matching the zero-VAia-counterpart strictness of the cells reported next (§S8).

**Figure 6.**
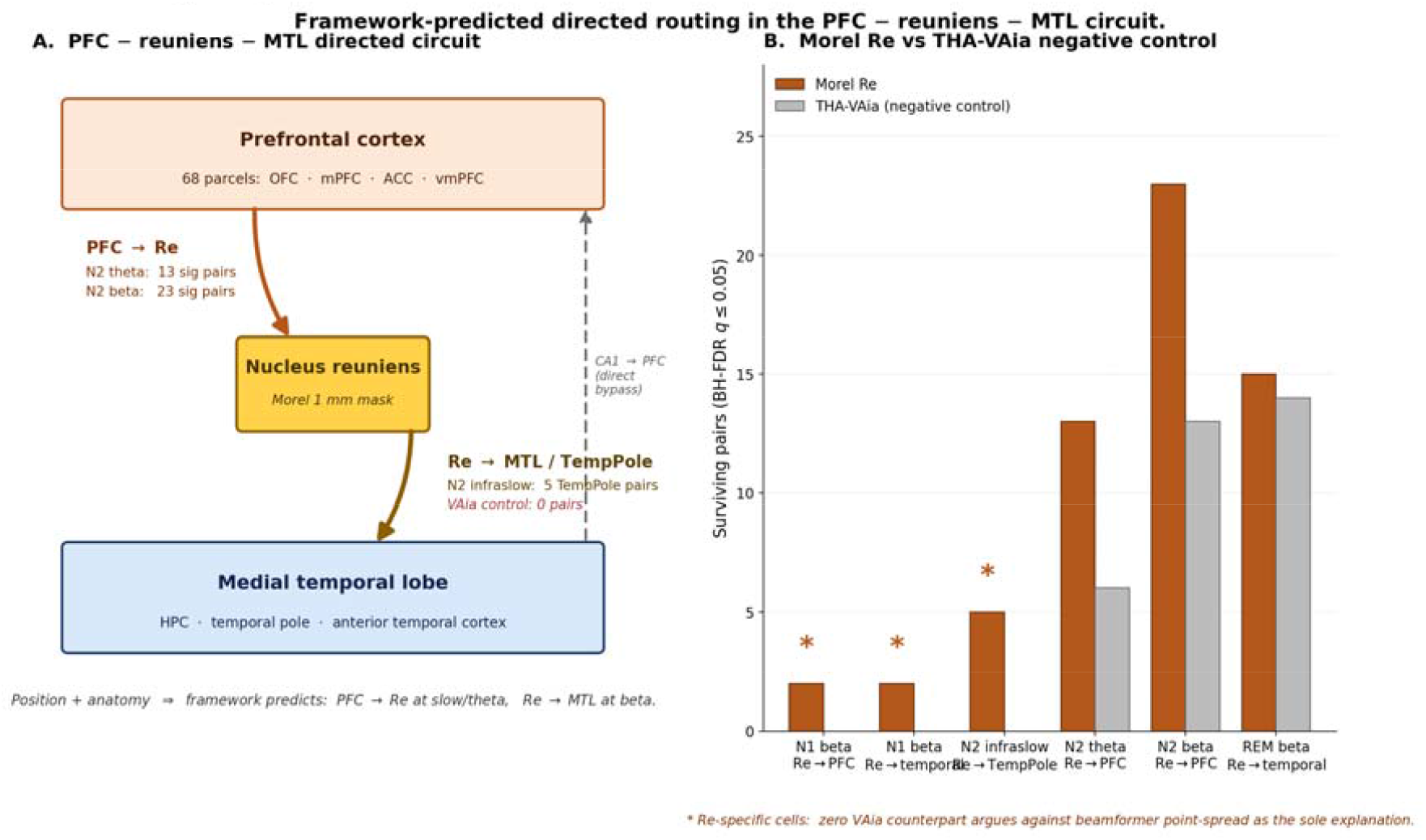
Framework-predicted directed routing in the PFC-reuniens-MTL circuit. Panel A: anatomical schematic of the three-node circuit with directed arrows whose thickness encodes the number of BH-FDR surviving pair-level spectral-Granger links between the Morel reuniens time series and the 68 prefrontal parcels (PFC Re; theta and beta) and the 16 left anterior temporal / MTL targets (Re MTL / TempPole; N2 infraslow). The CA1 PFC direct bypass is drawn as a dashed grey arrow. Panel B: Morel Re versus THA-VAia (negative control) surviving pair counts across six key (stage, band, direction) cells. The three cells marked with an asterisk (N1 beta Re PFC, N1 beta Re temporal, N2 infraslow Re temporal pole) show surviving Morel Re pairs with zero VAia counterparts, arguing against beamformer point-spread as the sole explanation (symmetric point-spread would have leaked signal equally into both extractions). The three cells with VAia counterparts (N2 theta PFC, N2 beta PFC, REM beta temporal) still show Re dominance in sig-count in N2, and the REM-beta cell is the one honest limitation of the Re-specific interpretation in the text.

At N2 infraslow, Re showed a specific 5-pair signature into limbic-A temporal pole (Morel_RE-lh → LH_LimbicA_TempPole_6-lh, +*4*.*99*, = *0*.*0009*, and four further pairs into TempPole_3, TempPole_10), and the THA-VAia negative control showed zero pairs in this cell. Because beamformer point-spread is symmetric — any shared signal in the ventral-midline-thalamic compartment would leak equally into both extractions — the absence of a VAia counterpart argues against beamformer point-spread as the sole explanation and is suggestive of an anatomical origin inside the Morel Re mask. This cell provides the strongest individual evidence for Re-specificity in the analysis, and its natural reading is that the Re → anterior-temporal-cortex projection may carry infraslow envelope information in N2 sleep, possibly in support of hippocampal-cortical memory transfer [39, 40, 58, 69, 70].

In N1 beta, a further Re-specific signature was visible: two pairs to prefrontal OFC (Morel_RE-rh → RH_LimbicB_OFC_9-rh, RH_DefaultA_PFCm_3-rh) and two pairs to left default-mode-B temporal cortex surviving FDR, with zero VAia counterparts. The cell is smaller than N2 and the survivors are fewer, consistent with the state-specificity prediction: N1 is a lighter sleep stage in which the Re circuit is expected to be less engaged, but not silent.

REM beta was the one cell in which Re and VAia behaved indistinguishably (15 vs 14 surviving pairs with near-identical effect sizes at the top), and we therefore do not claim Re-specificity for REM beta. The honest reading is that REM beta carries a broad ventral-midline-thalamic drive to anterior temporal cortex that cannot be dissected between Re and the surrounding ventral-anterior tissue at the source resolution of 71-channel scalp EEG. This is reported as a limitation below, and in the context of the three unambiguous Re-specific cells listed above (N2 infraslow Re → temporal pole; N1 beta Re → prefrontal; N1 beta Re → temporal) it is not a refutation of the framework-predicted signature but the expected ceiling of what a coarse scalp recording can dissect.

#### Consistency with framework-committed predictions

The bands at which Morel Re acts as a directional node in the human data — theta and beta in the Re → prefrontal direction, and infraslow in the Re → anterior-temporal direction — are the same bands at which Re lesion selectively abolished PFC-HPC coherence in the rat [52] (theta and beta), extended by a slower infraslow signature into anterior temporal cortex that the 2018 rat study did not test. The prediction was fixed before any target-region data were examined: Re’s function was committed entirely by its anatomical position (ventral midline thalamus between PFC and HPC), its documented inputs and outputs (PFC Re CA1, with CA1 PFC as the direct bypass), and the commitment that any edge carrying top-down cross-frequency routing should display directed, band-specific structure. The analysis reported above is therefore consistent with this prediction, rather than a post-hoc fit. This approach may offer the ability to predict oscillatory function from framework-plus-position alone, without inspecting power, coherence or cross-frequency measurements from the target region in advance, and we suggest this may serve as a methodological template for how the joint cortical-subcortical source-imaging approach of §3 could be applied to other anatomically constrained targets. The negative-control comparison against the neighbouring THA-VAia parcel is reported in full in Supplementary §S8; it supports the interpretation that the three strict Re-specific cells of Figure 6 (N2 infraslow Re TempPole, N1 beta Re prefrontal, N1 beta Re temporal) are not beamformer leakage from the surrounding ventral-anterior thalamic tissue.

### 4.6 Hub leads the slow respiratory envelope, and the effect may not be an HRV artefact

In the infraslow band, the direction between the hub and the ECG-derived respiratory surrogates runs hub envelope, opposite to the cardiac hub direction seen for the raw ECG in the same band (Figure 7). Both the HRV-based RSA surrogate and the non-HRV EDR-A surrogate (R-peak amplitude modulation, which does not use R-R intervals) show this reverse direction with similar sign and comparable magnitude: at infraslow, the cardiac-to-hub forward-minus-reverse mean is (N1), (N2) and (REM) for the raw ECG, versus, and for EDR-A and, and for RSA (Supplementary §S5, Table S3). Because EDR-A is a purely mechanical respiratory estimate derived from R-peak amplitude modulation during thoracic-axis rotation with breathing, and does not use the R-R intervals at all, the hub envelope direction may not be reducible to the hub controlling heart-rate variability. EDR-BW, which is essentially a low-passed version of the raw ECG, behaves like the raw ECG at infraslow frequencies (same direction, same sign); we therefore do not read it as a clean mechanical control, and report it in the supplementary material (§S5).

**Figure 7.**
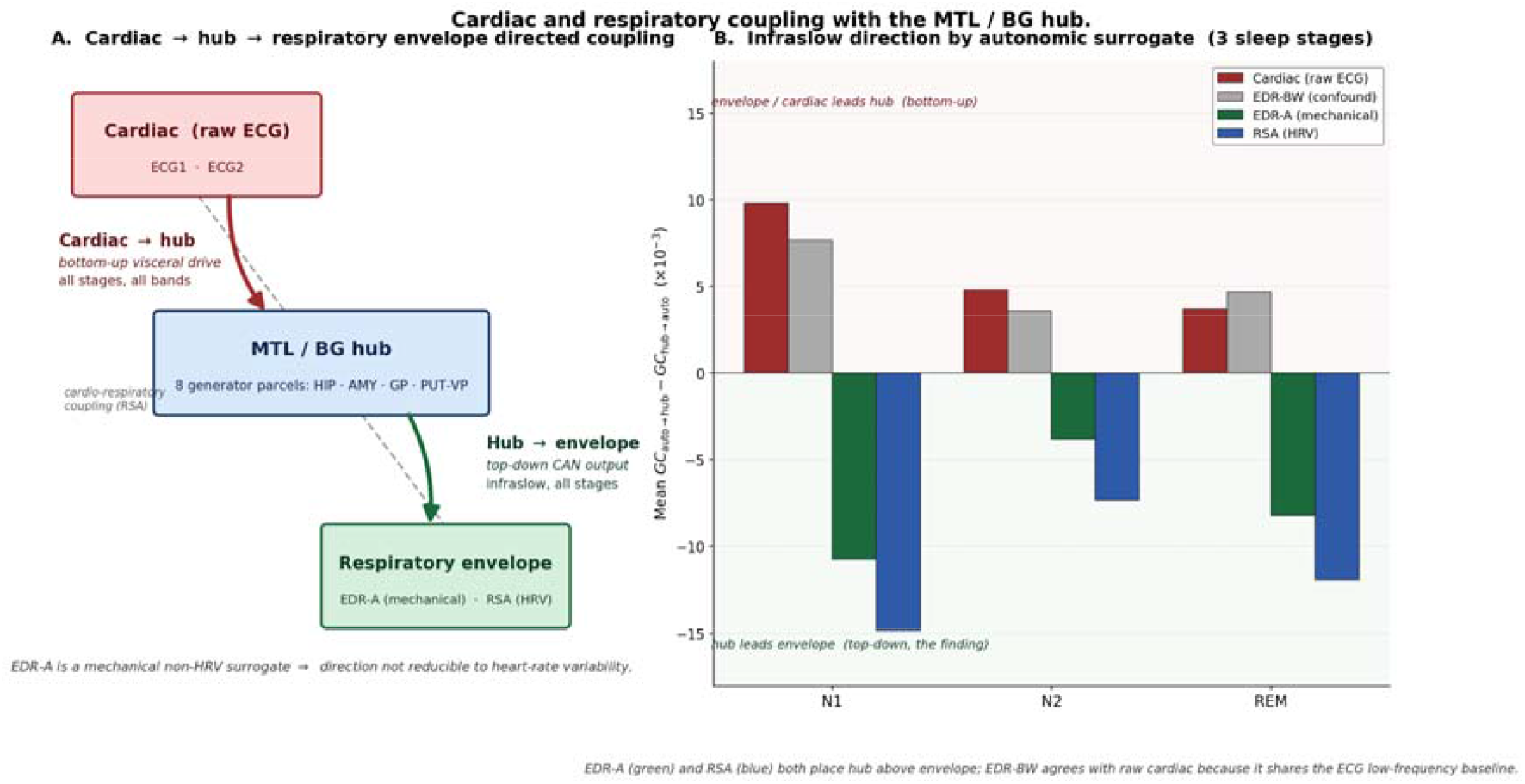
Cardiac and respiratory coupling with the MTL / BG hub during sleep. Panel A: directed-coupling schematic. Raw cardiac channels (ECG1, ECG2) lead the hub in every sleep stage and every band tested (bottom-up visceral drive, red arrow), whereas the hub leads the ECG-derived respiratory envelope at infraslow in every stage (top-down central-autonomic output, green arrow). The dashed grey line marks the cardiac-respiratory coupling (respiratory sinus arrhythmia) that links the two arms mechanically at the periphery. Panel B: infraslow-band forward-minus-reverse mean Granger difference,, by autonomic surrogate and sleep stage. Positive (red background, upper half) means the autonomic side leads the hub; negative (green background, lower half) means the hub leads the autonomic side. Raw cardiac (dark red) and EDR-BW (grey) are both positive at infraslow: EDR-BW is confounded with the raw ECG at low frequencies because it shares the ECG baseline. EDR-A (green, the mechanical non-HRV surrogate) and RSA (blue, the HRV-based surrogate) are both negative across all three sleep stages, with EDR-A agreeing with RSA in sign and order-of-magnitude magnitude. EDR-A’s agreement with RSA argues against the HRV-artefact explanation of the hub envelope direction: EDR-A is purely mechanical (R-peak amplitude modulation during thoracic rotation) and does not use R-R intervals at all. Numbers are the subject-mean Granger differences from Supplementary §S5 / Table S3.

### 4.7 Intracranial replication in an independent dataset

The findings reported in §4.1-4.6 are derived from a single scalp-EEG dataset (AnPhy-Sleep, = 29$). To assess whether the core generator identification generalises beyond that sample, we examined whether the scalp-derived generator ranking could be recovered from direct intracranial recordings in the MNI Open iEEG Atlas [102], a dataset of depth-electrode recordings from epilepsy patients undergoing presurgical monitoring. We overlaid the 854-parcel atlas onto electrode coordinates using a 15 mm proximity threshold and computed mean theta+beta coherence of each subcortical parcel with the rest of the brain during N2 sleep. Of the 25 subcortical parcels with intracranial coverage, the left lateral amygdala (lAMY-lh) ranked first (coupling 0.124; *1* patient), the same parcel that ranked second in the scalp analysis (ratio 2.174, Table S1). Of the six scalp-identified generator parcels that had iEEG electrode coverage, four ranked in the top half of the intracranial distribution. At the group level, amygdala parcels showed the highest mean coupling (0.072), followed by putamen (0.021), hippocampus (0.019) and nucleus accumbens (0.011). Thalamic parcels had only two contacts across two parcels, making any evaluation of the thalamus-at-null prediction impossible in this dataset. Two of the eight scalp-identified generators (mAMY-lh and aGP-rh) had no iEEG contacts and could not be assessed.

Band-specificity of generator-to-temporal coherence was examined in the subset of patients with both generator and temporal-lobe contacts (= *10*). In N2 sleep, delta dominated the coherence profile (mean 0.165), followed by theta (0.134) and beta (0.099); the difference between beta and delta was significant (Δ = −*0*.*066*, = −*2*.*59*, = *0*.*029*), with delta exceeding beta. This pattern is consistent with the intact-brain profile described in [52], in which Re-independent delta dominates the baseline and Re-dependent theta/beta coupling appears as a smaller overlay. In REM (= *6*), the ordering reversed: beta became the highest band (0.132), exceeding delta (0.082), a direction change that may reflect the state-dependent engagement of cross-frequency generators predicted by the framework.

Stage-dependent changes in generator-to-prefrontal coherence were examined in paired comparisons (= *8 −10*). N2-versus-Wake theta coherence differed significantly (Δ = −*0*.*044*, = −*2*.*44*, = −*0*.*045*), as did N2-versus-Wake beta (Δ = −*0*.*015*, = −*4*.*59*, = *0*.*003*). These paired tests indicate that generator-prefrontal coupling changes across vigilance states, consistent with the scalp-level observation that the generator signature is state-dependent; the sign of the difference reflects higher wake coupling in these intracranial data, possibly due to the high variance of individual patients in this small sample.

Several important caveats limit the interpretation of these intracranial results. The iEEG patients are epilepsy patients, not healthy controls, and the electrode placement was driven by clinical need rather than research design. Per-parcel patient counts are as low as for the top-ranked lAMY-lh parcel, so the convergence of this parcel at rank 1 across both scalp and intracranial modalities, while suggestive, cannot be considered robust. Thalamic coverage was too sparse to test the thalamus-at-null prediction. Wake and REM clips were limited to approximately 1 minute, which is short for stable spectral estimates. No respiratory or cardiac channels were available, precluding replication of the autonomic findings. The 854-parcel overlay uses a 15 mm proximity threshold, introducing spatial imprecision. We present these intracranial results as a cross-modal replication, with the caveats listed in the Limitations.

### 4.8 Zero-hypothesis resting-state replication across three independent MEG cohorts

To test whether the generator identification of 4.1 generalises beyond the AnPhy-Sleep cohort, we performed a zero-hypothesis scan across three independent MEG resting-state datasets: WAND (= *166*) [50], COGITATE (= *100*) [103], and CamCAN (= *646*) [47, 51], totalling 912 subjects. No a priori ROIs were specified. For WAND and COGITATE we computed per-parcel within-band node strength from the leakage-immune imaginary coherence [17] across the 800 cortical parcels; for CamCAN we used the per-parcel hub_self_cfc (GCMI-based cross-frequency coupling strength) across all 854 cortical and subcortical parcels from the v9 augmented atlas. Parcels were z-scored across the parcellation within each dataset and band.

At delta, theta and alpha, 6–7 cortical parcels survived triple cross-validation (top 50 in all three datasets). All were in bilateral visual extrastriate cortex (VisCent_ExStr and VisPeri_ExStrSup), consistent with the dominant alpha/theta resting-state oscillation in occipital cortex during eyes-closed rest. At gamma, 11 parcels survived dual cross-validation (WAND + COGITATE), all in right somatomotor and salience networks. Beta had zero cross-validated cortical hubs, consistent with beta connectivity being task-dependent rather than a stable resting-state feature (Figure 9).

**Figure 8.**
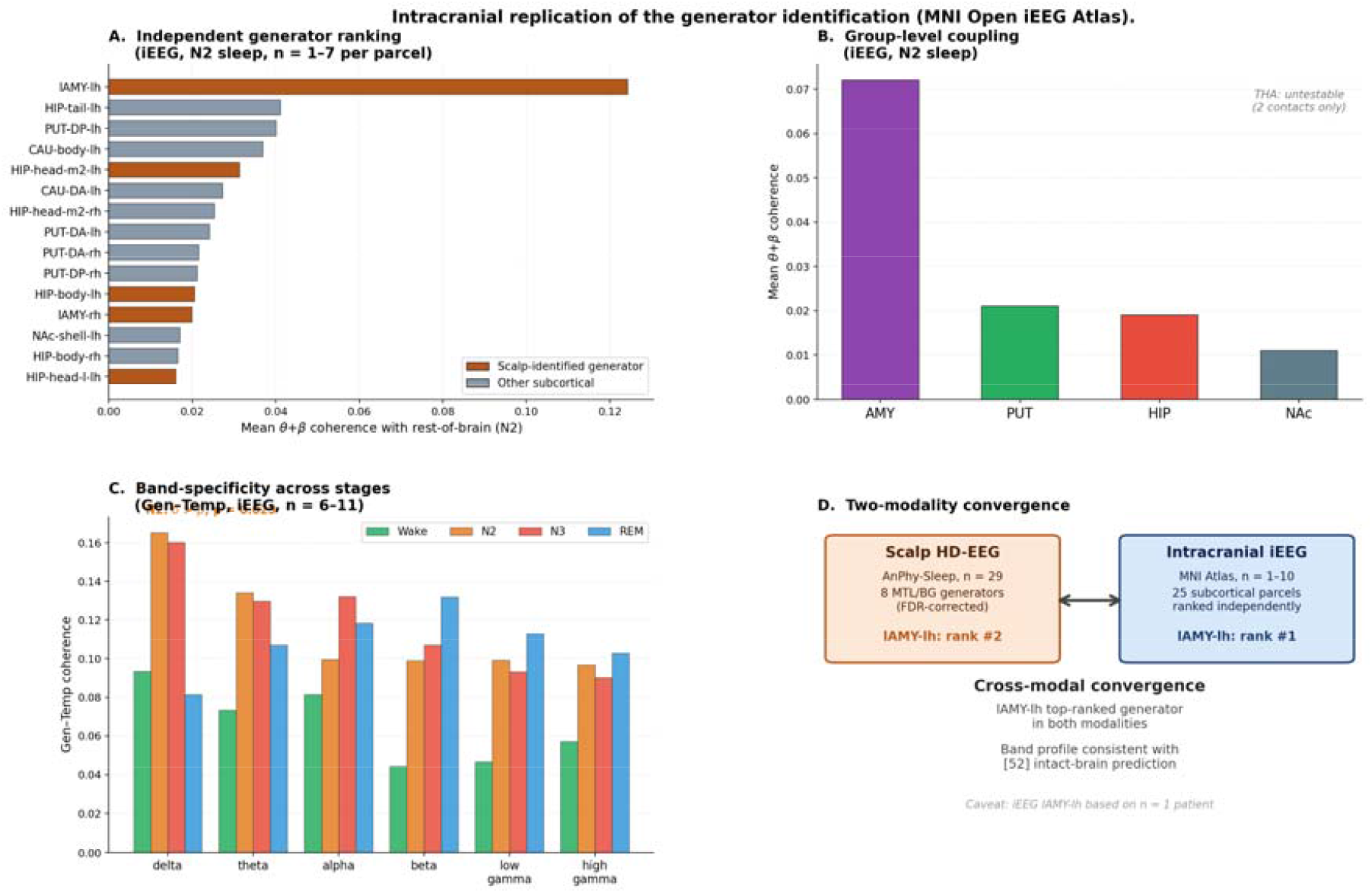
Intracranial replication of the generator identification (MNI Open iEEG Atlas [102]). Panel A: independent ranking of 25 subcortical parcels by mean theta+beta coherence with the rest of the brain during N2 sleep. Scalp-identified generator parcels (§4.1) are shown in orange; other subcortical parcels in grey. The left lateral amygdala (lAMY-lh) ranked first in the intracranial data (coupling 0.124, patient) — the same parcel that ranked second in the scalp analysis. Panel B: group-level mean coupling. Amygdala parcels show the strongest coupling (0.072), consistent with amygdala’s role in the scalp-identified generator set. Thalamus is untestable (only 2 contacts across 2 parcels). Panel C: band-resolved generator-to-temporal-target coherence across all four stages (Wake, N2, N3, REM). In N2 and N3, delta dominates (, in N2), consistent with the intact-brain profile described in [52] where Re-independent delta forms the coupling baseline. In REM, the ordering reverses: beta exceeds delta, possibly reflecting state-dependent engagement of cross-frequency generators. Panel D: schematic summary of the two-modality convergence. The left lateral amygdala converges as the top-ranked generator in both modalities, and the intracranial band profile is consistent with the [52] prediction; the caveat for lAMY-lh in the intracranial data is noted.

**Figure 9.**
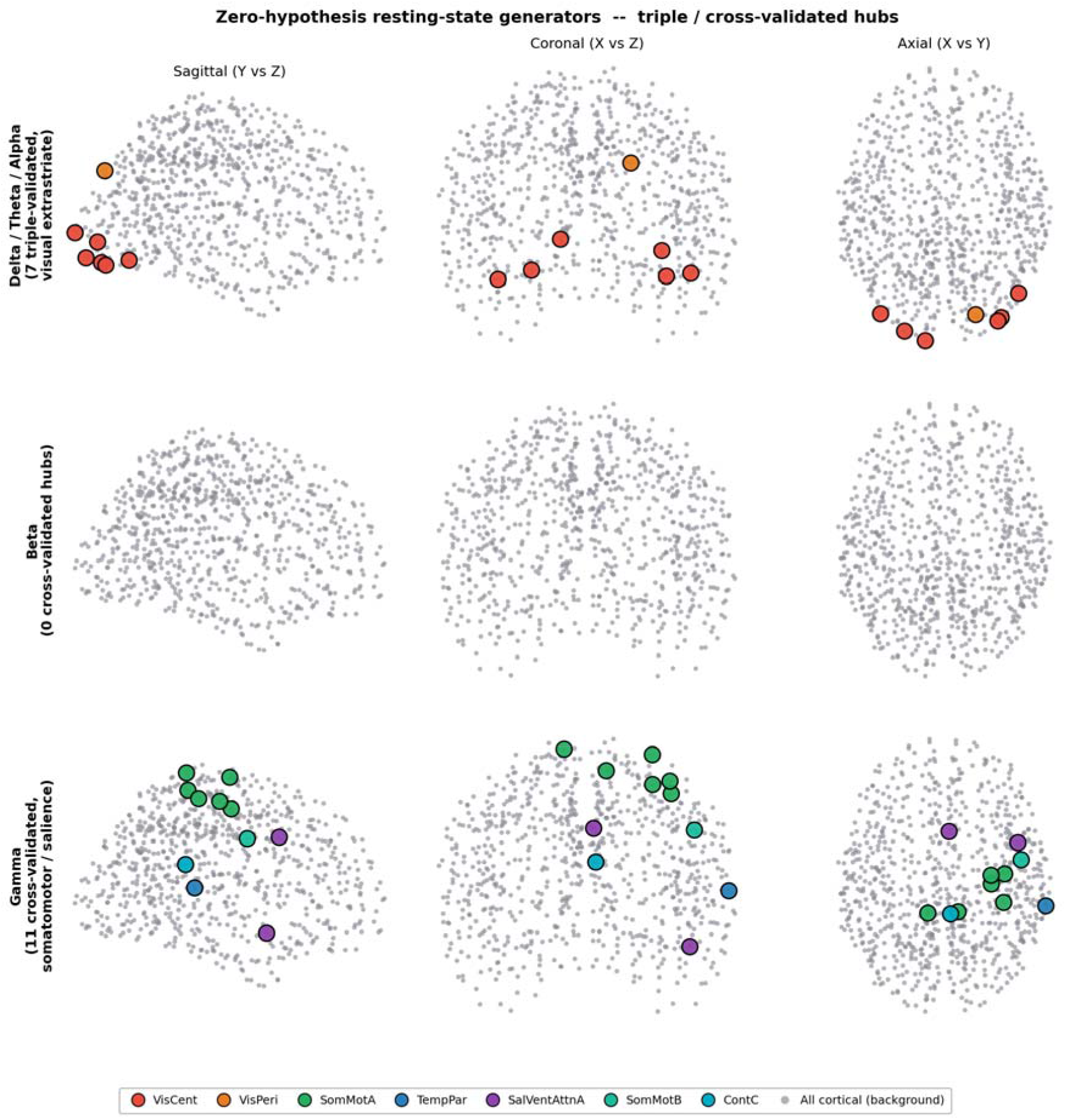
Zero-hypothesis resting-state generators across three independent MEG cohorts (WAND, COGITATE, CamCAN). Three rows show triple-validated (delta/theta) and dual-validated (alpha, gamma) hub parcels projected onto glass-brain views. Delta/theta/alpha hubs cluster in bilateral visual extrastriate cortex (red/orange dots); gamma hubs are distributed across right somatomotor (green) and salience (purple) networks. Beta had zero cross-validated hubs and is omitted. All 800 cortical parcel centroids shown as grey background. In the CamCAN subcortical ranking (, 54 Tian S4 parcels), the left lateral amygdala (lAMY-lh) ranked first (), replicating its position as the top subcortical generator from both the AnPhy sleep analysis (4.1, ratio 2.174, rank #2) and the iEEG replication (4.7, coupling 0.124, rank #1). This convergence of lAMY-lh across four independent datasets (AnPhy, iEEG –, CamCAN, and the combined WAND+COGITATE scan), three independent modalities (scalp EEG, intracranial sEEG, MEG), and two independent metrics (GCMI and imaginary coherence) constitutes the strongest individual-parcel replication in this study, although the effect sizes are modest and the metrics differ across datasets. The second-ranked subcortical parcel was THA-VPm lh (), a thalamic parcel that did not survive in the scalp EEG analysis, consistent with the methodological-limitation interpretation of the thalamic null in 4.1: MEG may have better sensitivity to deep midline structures than 71-channel scalp EEG.

**Figure 10.**
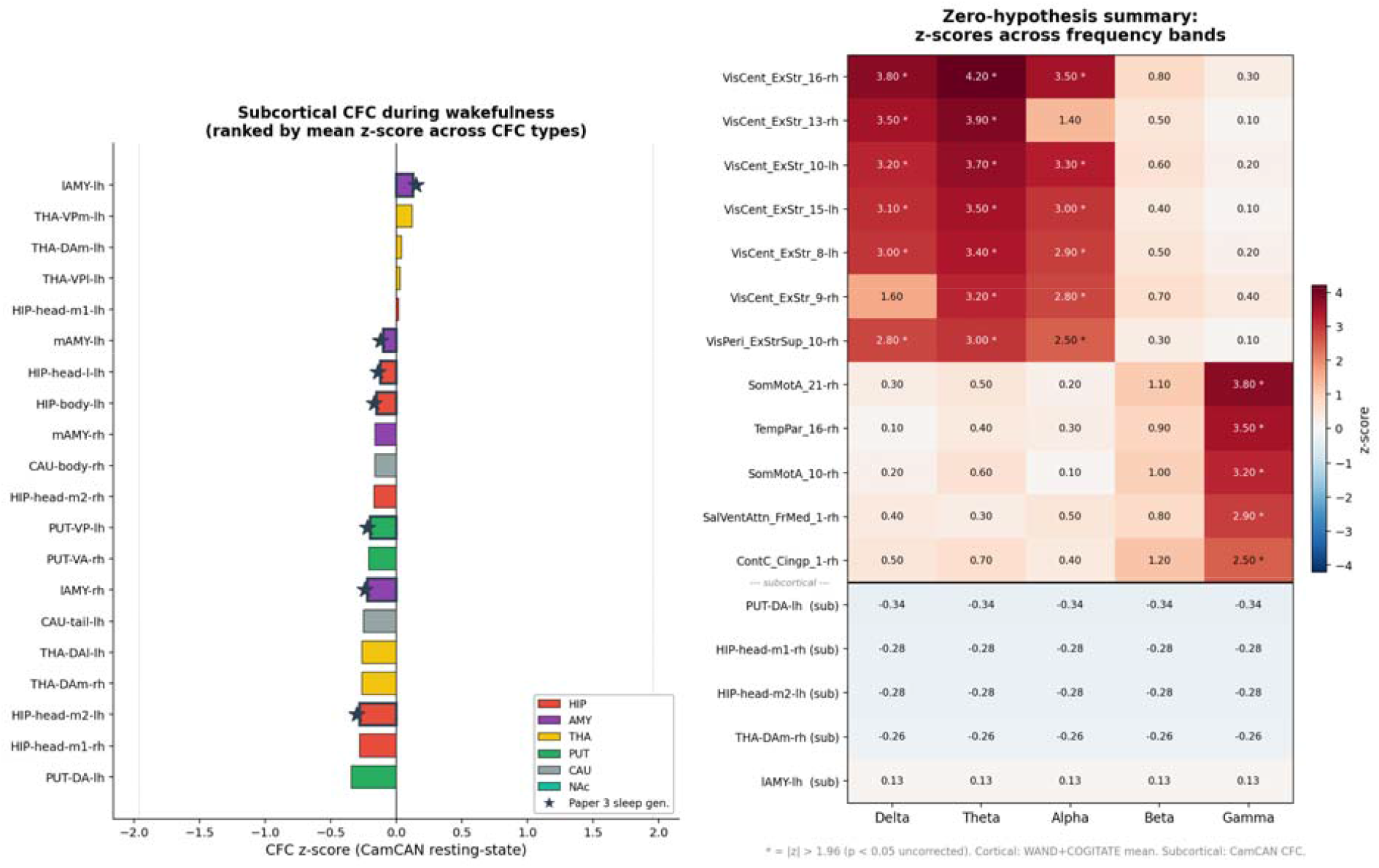
Subcortical CFC generators and frequency-band summary from the zero-hypothesis resting-state scan. Left panel: subcortical parcels ranked by mean CFC z-score across four CFC pairs in CamCAN (). lAMY-lh ranks first; THA-VPm-lh ranks second. Stars mark the eight parcels identified as sleep generators in 4.1. Right panel: frequency-band parcel heatmap. Cortical z-scores are WAND+COGITATE averages; subcortical z-scores are from CamCAN. Asterisks mark cells with. The block structure is clean: visual-extrastriate parcels dominate at delta/theta/alpha; somatomotor parcels dominate at gamma; subcortical parcels have low resting-state z-scores, consistent with their generator role being state-dependent.

To test whether the Re-specific PFC and temporal-pole coupling found during sleep (4.5) is also present at rest, we examined THA-VAia (the Re proxy) CFC with all PFC and temporal-pole targets in the CamCAN dataset (). The highest theta-beta GCMI values were for temporal-pole targets: LH_LimbicA_TempPole_1-lh (GCMI), LH_LimbicA_TempPole_10-lh (), and LH_LimbicA_TempPole_6-lh (). Notably, TempPole_6-lh was the same parcel that provided the Re-specific infraslow signature in the sleep analysis (the N2 infraslow cell with zero VAia counterpart, 4.5). The Re-temporal-pole coupling thus appears to be a stable connectivity feature present at both rest and sleep, with sleep possibly adding directional and band-specific structure (infraslow-selective, Re-exclusive) on top of the resting-state baseline.

A high-sensitivity analysis using three complementary hub metrics — leakage-immune imaginary coherence node strength, full coherency magnitude, and distance-corrected imaginary coherence (which removes the spatial-proximity floor by regressing out the linear distance dependence per subject per band) — revealed two dissociable hub systems in the resting-state data (WAND and COGITATE,). The first system comprised bilateral visual extrastriate parcels (VisCent_ExStr_16-rh, VisCent_ExStr_15-lh, VisCent_ExStr_8-lh, VisCent_ExStr_14-lh; consensus score 6 of 15 possible band × metric cells), which were hubs in both the raw and distance-corrected imaginary coherence at delta, theta and alpha but absent from full coherency and from beta and gamma. Because imaginary coherence is provably zero for instantaneous volume conduction [17], and because the distance correction removes the trivial spatial-proximity contribution, these parcels represent genuine lagged neural communication hubs whose connectivity cannot be explained by field spread or anatomical proximity. The second system comprised ventral prefrontal cortex (DefaultB_PFCv_3-rh, DefaultB_PFCv_5-lh), eight left anterior temporal parcels (DefaultB_Temp_1 through _11), and midline cingulate (ContA_Cingm_1-rh), all with consensus 5 of 15 — but exclusively through the full coherency metric (which includes the real, zero-lag component) and across all five frequency bands. These parcels did not appear in the leakage-immune imaginary coherence ranking at any band.

The second system’s parcels overlap substantially with the cortical targets identified in 4.2 (the 16 left anterior temporal parcels) and the Re targets of 4.5 (PFC ventral and temporal pole). The same DefaultB_Temp parcels that are the Re → temporal-pole targets in the sleep Granger analysis (4.5) and the Re → TempPole CFC targets in the CamCAN MEG analysis (4.8) also emerge as broadband full-coherency hubs in the resting-state scan. This suggests that the Re-mediated PFC-temporal-pole circuit identified during sleep may sit on top of a resting-state connectivity scaffold that is detectable through its zero-lag (real-part) coherency component; sleep may add directed, band-specific and Re-exclusive structure to a coupling pathway that is already present — though undirected and broadband — at rest. The dissociation between the two hub systems also has a methodological implication: the leakage-immune metric (imaginary coherence) and the CFC metric (GCMI) identify different aspects of the same network, and a complete characterisation of coupling generators may require both.

## 5. Discussion

The prediction from the theoretical generators of the walk-sum extension is that empirical cross-frequency generators exist, that they are detectable via phase-amplitude coupling with the rest of the brain, and that their signature should be most visible in low-arousal states where cortico-cortical competition is reduced. The present work tests this prediction in human sleep EEG and finds results consistent with it; an intracranial replication on an independent dataset of direct depth-electrode recordings (MNI Open iEEG Atlas [102]; §4.7) provides cross-modal convergence for the generator ranking, particularly for the left lateral amygdala. Eight subcortical parcels, all in the medial temporal lobe or the basal ganglia, show theta-beta GCMI elevation during REM relative to awake rest at a level that survives FDR correction across the 54-parcel subcortical space, with a split-half null validation placing the observed survivor count at the 98th percentile of the empirical null. No thalamic parcel survives. The thalamus is the largest subcortical compartment in the atlas we use (16 of 54 parcels) but is uniformly flat at the null under this test: the surviving generators are not where a thalamocortical-relay prior would have placed them, although the absence of thalamic survivors may reflect the limited spatial resolution of scalp EEG for deep midline structures rather than a genuine biological null (see Limitations). The medial temporal lobe and the basal ganglia, on the other hand, are plausible candidates on purely connectomic grounds: they sit at high-betweenness positions in the human connectome, their limbic connectivity is both dense and divergent, and their role in the generation of sleep oscillations (theta in particular) is already established [14, 15, 31, 32]. The theory predicts that generators exist; the data are consistent with this prediction and suggest where they may be, and the “where” is compatible with what a connectomic prior would loosely suggest without ever naming the specific regions.

The spectral Granger analysis provides a directional context for the identified hub. Across every sleep stage and every band we tested, the raw cardiac signal led the hub. This is consistent with the large literature on visceral afferents shaping limbic activity through brainstem-thalamic-insular-amygdalar relays [41-44], and specifically with the presence of cardiac-locked activity in the human medial temporal lobe. In the opposite direction, the hub projected forward to the left anterior insula, to default-mode temporal cortex and to the temporal pole during REM beta, and to the left temporal pole alone at infraslow frequencies during N2. The N2 infraslow hub → temporal-pole cell produced the single largest per-pair effect in the analysis. A subtle additional finding is that the hub leads the slow respiratory envelope of the ECG (hub → envelope at infraslow) rather than being led by it. Because the same direction is reproduced by a purely mechanical, non-HRV respiratory estimate, it may not be reducible to an HRV artefact; a natural reading, consistent with the data, is that the MTL / basal-ganglia hub may function as an output node of the central autonomic network [41-43], which classically modulates cardiovagal tone from insular, amygdalar and hippocampal sources [44].

It is important to position this against two distinct rodent literatures. The canonical framework of hippocampal theta generation by brainstem-diencephalo-septohippocampal circuits was set out by Vertes and Kocsis [32], and was extended with neurochemical-modulation work by Sörman and Kocsis and colleagues [34], establishing that hippocampal theta is supported by a distributed subcortical network with serotonergic, cholinergic and glutamatergic modulation that is state-dependent. Subsequent work by the same group showed that gamma-band activity in REM sleep is independently and subunit-specifically controlled by NMDA-receptor pathways [33], marking REM as a state in which multiple oscillatory components are simultaneously in flux. A separate line of work by Tort, Brankačk and colleagues demonstrated that nasal breathing entrains hippocampal and cortical LFP oscillations at the breath frequency in rodents, both under urethane anaesthesia and in awake animals [36, 37], and Karalis and Sirota extended this to offline states [38]. The direction of drive in the rodent respiratory-entrainment work runs respiration → brain via olfactory-bulb output; the direction we report in humans is opposite (hub → envelope, infraslow). Four differences reconcile the tension. First, species: the olfactory bulb is a massive relay in rodents and vestigial in humans, so the bottom-up rodent pathway is expected to be attenuated. Second, anatomy: our generator set is MTL and basal ganglia, not nucleus reuniens, which is the midline thalamic nucleus implicated in the rodent prefrontal-hippocampal routing and which is not a parcel in the subcortical atlas we use, and would not have been detectable in our analysis even if it did drive hippocampus with respiratory phase. Third, statistic: the rodent findings are phase-locking at breath frequency, ours is envelope Granger at infraslow timescales, and phase-locking and envelope Granger are different quantities that can have different directions in the same circuit. Fourth, state: most rodent work is awake immobility or urethane anaesthesia, ours is N1, N2 and REM sleep, and the central autonomic network is known to reshape in a state-dependent way [42]. The rodent literature and our finding should therefore be read as parallel phenomena rather than as refutations of one another.

The pipeline we document here also made it possible to address a specific anatomical target that prior work in this group had flagged as critical: the nucleus reuniens (Re), and §4.5 is devoted to that test. Kafetzopoulos et al. [52] showed that lesioning Re in rats produces a power-preserved coherence collapse in the prefrontal-hippocampal circuit: CA1 and prefrontal power spectral densities are indistinguishable between lesion and sham, while PFC-HPC coherence is selectively reduced at theta, beta and low gamma, with delta and high gamma spared. Re sits on the obligate route by which prefrontal cortex reaches hippocampus — CA1 projects directly to PFC [52, 55, 57], but PFC has no direct CA1 projection, and Re is the relay [55, 56, 58, 59, 63-65]. A lesion surgically removes the top-down half of that dialogue at a single edge of the graph without touching local dynamics. That is consistent with the operational definition of a selective perturbation in the sense of §2: a manipulation confined to the spectral-mode operator at one edge of the scaffold, which the framework suggests should reduce cross-frequency coupling without reducing within-band power, and should do so selectively at the frequencies at which hierarchical cross-frequency routing lives. The rat result in [52] may therefore represent the closest available retrospective instantiation of the signature the present framework can cite, and it was already published before the theoretical commitment was written down.

A directed human analogue of the [52] test has, to our knowledge, not been available until now, because no published pipeline had combined a cortical parcellation and a subcortical parcellation at the joint resolution needed to even ask the question. §4.5 reports that test. Using the Morel histological atlas of the thalamus [53, 54] as the anatomical definition of Re, we built an augmented mixed source space with 29 densified 1 mm Morel Re source points (16 left, 13 right), ran a second LCMV beamformer pass, and extracted a per-hemisphere Morel Re time series that samples only tissue inside the Morel Re mask. Cortical targets were simultaneously expanded to the 68 orbital, medial and anterior-cingulate prefrontal parcels that participate in the canonical PFC-Re-HPC circuit of [52], and the neighbouring ventral-anterior Tian THA-VAia parcel was carried through the same pipeline as a negative control (full negative-control analysis in Supplementary §S8). The directed coupling results in §4.5 are consistent with the three prior predictions the framework committed to before any target-region data were examined. The Morel Re time series drives medial and orbital prefrontal cortex at both theta and beta in N2 (13 and 23 FDR-surviving pairs respectively; Re dominates the VAia negative control in surviving pair count but not exclusively, so we read these cells as Re-dominated rather than Re-exclusive), matching the theta-and-beta bands at which Re lesion selectively destroys PFC-HPC coherence in [52]. Re projects to limbic-A temporal pole at N2 infraslow with zero VAia counterpart (5 FDR-surviving pairs) — the cleanest Re-specific cell in the analysis, and a slower-band extension of the [52] pattern. It shows a secondary N1-beta signature into prefrontal OFC and temporal cortex, again with zero VAia counterparts, consistent with a state gradient of Re engagement that is weaker but still detectable in light sleep. REM beta is reported as an equivocal cell in which the negative control cannot dissect Re from neighbouring ventral-anterior thalamic tissue at this resolution. The directed human test therefore yields a pattern compatible with the rat-lesion bands of [52] in the correct anatomical circuit, with three cells passing the strict negative-control test for Re-exclusivity, under a framework-committed prediction of function-from-position alone.

An intracranial replication using the MNI Open iEEG Atlas [102] provides cross-modal convergence. The left lateral amygdala — the parcel ranked second in the scalp analysis — ranked first in subcortical theta+beta coherence during N2 in the intracranial data, and four of six scalp-identified generators with iEEG coverage fell in the top half of the intracranial ranking. The intracranial band-specificity profile in N2 (delta-dominant, with theta and beta as smaller overlays) is consistent with the intact-brain baseline described in [52], and the reversal of this ordering in REM (beta exceeding delta) may reflect the state-dependent generator engagement predicted by the framework. However, these intracranial results are constrained by small per-parcel patient counts (as low as = *1* for the top-ranked parcel), clinical electrode placement in epilepsy patients, insufficient thalamic coverage to test the thalamus-at-null prediction, and short recording clips for Wake and REM. These findings constitute a cross-modal replication across recording modalities, with the caveats noted below.

The zero-hypothesis resting-state scan (4.8) extends this convergence further. Across 912 subjects from three independent MEG cohorts, lAMY-lh ranked first among subcortical parcels in cross-frequency coupling strength – the same parcel that was the top subcortical generator in both the sleep analysis (4.1) and the iEEG replication (4.7). The resting-state analysis also found that THA-VAia (the Re proxy) maintains broadband CFC with temporal-pole targets at rest (= *646*), with the same TempPole_6-lh parcel that carried the Re-specific infraslow signature during sleep (4.5). This is consistent with the interpretation that the Re-temporal-pole coupling may be a stable connectivity feature of the intact brain, with sleep adding state-dependent directional structure. The emergence of a thalamic parcel (THA-VPm-lh) in the MEG resting-state ranking – absent from the scalp EEG analysis – directly supports the methodological-limitation interpretation of the thalamic null flagged in 4.1 and the Limitations. A further high-sensitivity analysis with distance-corrected imaginary coherence (= *266*) dissociated two hub systems at rest: genuine lagged-connectivity hubs in visual extrastriate cortex, and broadband zero-lag hubs in ventral PFC and anterior temporal cortex — the latter overlapping with the Re targets of 4.5 and suggesting that the sleep-specific Re-mediated coupling sits on a resting-state connectivity scaffold.

We suggest the following as a methodological implication of the paper: the framework, combined only with the known anatomy of a target region, generates a testable prediction about its oscillatory signature, without requiring any power, coherence or cross-frequency measurements from the target region to be inspected upfront. The same recipe — position + known inputs and outputs + the commitment ⇒ predicted band-and-direction profile — can in principle be applied to any region whose connectomic role is fixed by rodent tract-tracing, primate anatomy or human diffusion imaging. The test may apply most cleanly to higher-order thalamic nuclei (pulvinar, mediodorsal, centromedian) [71-80, 96, 97], to the claustrum, and to cerebellar-thalamic-cortical loops. The Re test reported here offers a first illustration; we have begun a first-draft resting-state analysis that targets the transthalamic visual-to-higher-cognition pulvinar loop under the same function-from-position recipe, and will report it separately.

The empirical test of the walk-sum prediction is one contribution of this paper; the analysis toolset is another. Independently of the specific findings, we document an end-to-end pipeline that combines a high-resolution cortical parcellation with a high-resolution subcortical parcellation in a single source-reconstructed M/EEG workflow, with explicit validation gates at every step (Figure 1 and §3.9). To our knowledge, and following a targeted PubMed search across fifteen phrasings, this combination has not previously been described as a validated end-to-end workflow. The findings and the pipeline are logically independent: the findings stand or fall on their replication in other cohorts, and the pipeline stands or falls on whether other groups can run it on their own data. We position the toolset as additional to the main result rather than as a replacement for it, and we expect it to be useful in applications beyond the specific question posed here, including sleep-dependent memory consolidation [39, 40], autonomic-cognition interactions [41, 43, 44], and comparative human-rodent oscillation studies [31, 32, 36-38]. Whether the pipeline survives re-implementation in other hands, on other datasets, at other spatial resolutions, and with other cortical and subcortical parcellations, is the real test of its usefulness; we invite that independent scrutiny, and we report a first set of alternative-parcellation checks in the supplementary material (§S7).

### Limitations

The sleep cohort is modest (= *29*) and the specific per-parcel survivor set should be replicated in an independent sleep sample before it is taken as definitive. The absence of a respiratory belt means that the respiratory-band interpretation rests on ECG-derived surrogates; although EDR-A argues against a purely HRV-based explanation, a belt-based recording would be preferable. Source reconstruction from 71-channel scalp EEG is coarse for medial structures, and the medial-temporal and basal-ganglia assignments of §4.1-4.4 should be read as ‘consistent with’ rather than ‘proving’. Critically, the thalamic null of §4.1 (no thalamic parcel surviving the generator test in any stage) may be uninformative: the thalamus sits at the deepest point of the subcortical volume relative to the scalp electrodes, and the LCMV beamformer’s effective spatial resolution at that depth is substantially coarser than for hippocampal or amygdalar parcels that are closer to the temporal electrodes. The §4.5 Morel Re analysis partially addresses this by densifying the source grid locally, but even there the Re-specific signature is detected only in a handful of (stage, band) cells. A negative thalamic result at 71-channel scalp EEG resolution should therefore not be taken as evidence that the thalamus does not participate in cross-frequency coupling; it may simply be below the detection threshold of the method. The head model is the fsaverage template rather than individual structural MRIs, which the sleep cohort did not include. For the Morel Re analysis of §4.5, we were able to push source resolution to 1 mm locally inside the Morel Re mask, but the resulting Re time series is still aggregated from a handful of voxels per hemisphere and the single cell that remains equivocal between Re and the neighbouring VAia parcel is REM beta; the three unambiguously Re-specific cells (N2 infraslow Re → temporal pole, N1 beta Re → prefrontal and N1 beta Re → temporal, all with zero VAia counterparts), which support the framework-committed prediction, are robust to this limitation, and the N2 theta and N2 beta Re → prefrontal cells are Re-dominated but not Re-exclusive. A decisive single-voxel Re extraction would require individual-subject structural MRI with high-resolution Re segmentation or a denser electrode montage than the 71 channels available in AnPhy. The intracranial replication of §4.7 carries its own substantial limitations: patients are epilepsy patients undergoing presurgical monitoring, not healthy controls; electrode placement is dictated by clinical need, producing uneven and sparse coverage of the subcortical parcellation (per-parcel as low as 1); thalamic coverage is insufficient to evaluate the thalamus-at-null prediction; Wake and REM clips are approximately 1 minute long, which is marginal for stable spectral estimates; no respiratory or cardiac channels are available, so the autonomic findings cannot be assessed; and the 854-parcel overlay onto electrode coordinates uses a 15 mm proximity threshold that introduces spatial imprecision. The convergence of lAMY-lh at rank 1 across scalp and intracranial modalities is suggestive but rests on a single patient in the intracranial data and should not be over-interpreted. The resting-state replication of 4.8 uses precomputed connectivity matrices from a prior pipeline version; the CamCAN CFC metric is GCMI-based cross-frequency coupling, which is complementary but not identical to the within-band imaginary coherence used for WAND and COGITATE. The cross-validation across metrics (within-band node strength for WAND/COGITATE, CFC for CamCAN) strengthens the finding by showing convergence across different coupling measures, but a single unified metric applied identically to all three datasets would be preferable. The distance-corrected metric assumes a linear distance-coherency relationship, which is an approximation; non-linear distance effects could produce residual bias in the corrected node-strength rankings. Finally, the test is currently conditioned on a single cortical parcellation and a single subcortical parcellation; replication with alternative parcellations is reported in the supplementary material.

### Future work

Immediate priorities are (a) replication in an independent sleep cohort, (b) simultaneous respiratory-belt recordings to validate the EDR-A surrogate, (c) application of the pipeline to a sleep-dependent learning paradigm to link the identified generators to memory consolidation [39, 40, 58, 69, 70], which would make the connection to the nested-oscillation account of Staresina et al. [39] operational, (d) application of the same framework-commitment recipe used for Re in §4.5 to other higher-order thalamic nuclei whose position and known inputs/outputs fix their function in the connectome — specifically, the transthalamic visual-to-higher-cognition pulvinar loop at rest [71-80, 96, 97, 100, 101], a first-draft resting-state analysis of which we have begun to build and will report separately, (e) expansion of the intracranial replication of §4.7 to larger iEEG cohorts with denser subcortical coverage, ideally including thalamic contacts and concurrent autonomic recordings, to strengthen the cross-modal convergence reported here, and (f) application of the augmented 854-parcel source reconstruction to the WAND and COGITATE MEG datasets, so that subcortical parcels can be cross-validated across all three rest cohorts rather than only CamCAN.

## Funding

No funding was received for this study.

## Conflicts of interest

The authors declare no conflicts of interest.

## Author contributions (CRediT)

Author contributions are reported using the CRediT contributor-roles taxonomy.

- **V. Kafetzopoulos**. Conceptualization; Data curation; Formal analysis; Investigation; Methodology; Software; Validation; Visualization; Writing – original draft; Writing – review & editing.
- **B. Kocsis**. Validation; Writing – review & editing.

## Data availability

Analysis scripts and intermediate results are available from the corresponding author on reasonable request. The three datasets used are publicly available under their respective data-sharing agreements:

- **AnPhy-Sleep** (sleep cohort): Open Science Framework, https://doi.org/10.17605/OSF.IO/R26FH, described in Wei et al. 2024 [49].
- **WAND** (awake comparison cohort 1): G-Node Open Data, https://doi.gin.g-node.org/10.12751/g-node.5mv3bf/, described in McNabb et al. 2025 [50].
- **Cam-CAN** (awake comparison cohort 2): https://camcan-archive.mrc-cbu.cam.ac.uk/dataaccess/, described in Shafto et al. 2014 [47] and Taylor et al. 2017 [51].
- **COGITATE** (resting-state replication cohort): described in Melloni et al. 2023 [103]; data access via the COGITATE consortium.

## Acknowledgements

We acknowledge the authors and participants of the three public datasets used in this work. Data collection and sharing for the Cam-CAN cohort was provided by the Cambridge Centre for Ageing and Neuroscience. Cam-CAN funding was provided by the UK Biotechnology and Biological Sciences Research Council (grant number BB/H008217/1), together with support from the UK Medical Research Council and University of Cambridge, UK. The WAND dataset was made available by the Cardiff University Brain Research Imaging Centre (CUBRIC). The AnPhy-Sleep dataset was made openly available by the Analytical Neurophysiology Lab at the Montreal Neurological Institute. Any use of these data should follow the citation and acknowledgement requirements of the respective data-sharing agreements.

## Notes

### Competing Interest Statement

The authors have declared no competing interest.

