## Supplementarry Text for "Walk-sum theoretical generators predict medial-temporal coupling driven by empirical cross-frequency generators: a novel joint cortical-subcortical approach in human sleep EEG"

**V. Kafetzopoulos**${}^{1}$**, B. Kocsis**${}^{2}$

${}^{1}$*Department of Psychiatry, Medical School, University of Cyprus, Nicosia, Cyprus*

${}^{2}$*Department of Psychiatry, Beth Israel Deaconess Medical Center, Harvard Medical School, Boston, MA, USA*

This document contains the full theoretical derivation, a review of alternative phase-amplitude coupling metrics held as a reviewer reserve, the full per-parcel tables of the generator analysis, a split-half null validation, a breakdown of the spectral-Granger results by stage, band and direction pair with the four-group autonomic rollup, alternative-parcellation robustness checks, and quality-control details.

### S1. Full theoretical derivation

#### S1.1 Walk-sum resolvent

Consider a discrete system of $N$ regions, each carrying a scalar signal, coupled through an $N\times N$ weighted adjacency matrix $C$. Let the intrinsic regional dynamics be approximated by a linear filter $H_{0}(\omega)$ at each node and let the inter-regional delay be $T$. The steady-state response of the system to a stimulus $s(\omega)$ satisfies

$$X(\omega)=H_{0}(\omega) X(\omega)+H_{0}(\omega) e^{i\omega T} C X(\omega)+H_{0}(\omega) s(\omega).$$

Collecting terms,

$$X(\omega)=H_{0}(\omega) [I-e^{i\omega T}C]^{-1} s(\omega),$$

and the response-matrix factor on the right of (S2) is the dressed resolvent

$$H(\omega)=[I-e^{i\omega T}C]^{-1}=\sum_{k=0}^{\infty} (e^{i\omega T}C)^{k}.$$

The Neumann series in (S3) converges whenever $\rho(C)<1$. Each term $k$ is a sum of matrix products of $k$ copies of $C$, so $(H)_{ij}$ is a sum over all directed walks of length $k$ from $j$ to $i$, weighted by the product of edge weights along the walk and phase-delayed by $kT$. This is the formal meaning of the phrase “walk-sum on the connectome”.

#### S1.2 Why scalar is insufficient for multi-band dynamics

The scalar resolvent assigns a single complex number to each ($\omega$, edge) pair. This is the correct operator only when regional dynamics can be summarised by a single spectral mode. In the cortex and subcortex, regional dynamics empirically cannot be so summarised: there are at minimum five well-characterised bands (delta, theta, alpha, beta, gamma); their amplitudes modulate each other through cross-frequency coupling; and the phase of one mode in one region carries statistical information about the amplitude of a different mode in a different region [11, 12, 13, 14, 15]. Insisting on a scalar resolvent forces one either to collapse across bands, which loses information, or to stack them additively, which forbids interactions of the form “the amplitude of the fast mode is modulated by the phase of the slow mode”: the cornerstone of cross-frequency coupling.

#### S1.3 Replacing the scalar with a spectral-mode operator

Let the regional state at frequency $\omega$ now be a vector in a band space $X\in\mathbb{C}^{N\times M}$, where $M$ is the number of bands. At each hop the propagator is no longer a scalar times $C_{ij}$, but an operator acting on the band space. Denote this operator at edge $e$ as $G_{e}\in GL(M,\mathbb{C})$. The band-space propagator of a single hop is

$$\rho(e)=G_{e} e^{i\omega T} C_{ij(e)}.$$

For a walk $\gamma=e_{1}e_{2}\cdots e_{k}$ of length $k$, the band-space propagator is the path-ordered product of hop operators in the order the hops are traversed,

$$\rho(\gamma)=\mathcal{P}\prod_{\ell=1}^{k} \rho(e_{\ell})=G_{e_{k}}\cdots G_{e_{2}} G_{e_{1}} e^{ik\omega T} \prod_{\ell} C_{ij(e_{\ell})}.$$

The $G_{e}$ do not in general commute, so $\rho(\gamma)$ depends on the order in which hops are taken. The dressed-resolvent analogue is the sum over all walks of all lengths,

$$\mathcal{H}(\omega)=\sum_{\gamma} \rho(\gamma)=\mathcal{P}\exp\left[ \int_{\text{hops}} G_{a(\ell)} d\ell\right],$$

with the second equality holding as a formal path-integral continuum limit. If the $G_{e}$ can be exponentiated as elements of a Lie group that preserves total band-space power and acts by unitaries, the natural choice is $SU(M)$. The generators of $SU(M)$ form a basis for traceless anti-Hermitian matrices and correspond to the spectral-mode mixings permitted by the theory. The non-abelian nature of $SU(M\geq2)$ means that the ordering of hops is informative: this is the formal statement of the informal observation that “the brain is more than a weighted graph”.

#### S1.4 From the operator to empirical observables

In the scalar theory the natural two-point function is the cross-spectrum

$$S_{ij}(f)=\mathbb{E}[X_{i}(f) X_{j}^{*}(f)].$$

In the non-abelian theory, the lowest-order gauge-invariant observable sensitive to non-commutativity is the three-point cross-bispectrum

$$B_{ijk}(f_{1},f_{2})=\mathbb{E}[X_{i}(f_{1}) X_{j}(f_{2}) X_{k}^{*}(f_{1}+f_{2})].$$

The cross-bispectrum is a direct analogue of a three-point correlator in non-abelian gauge theories, and it carries information that the two-point function cannot, namely the phase of the slow mode at site $i$, the amplitude of the fast mode at site $j$, and the complex phase relationship between them. The magnitude normalisation is bicoherence (equation 5 in the main text), and the antisymmetric combination of Chella et al. [16] is provably insensitive to instantaneous mixing artefacts:

$$b_{anti}(f_{1},f_{2})=\frac{\left| \mathrm{Im}B(f_{1},f_{2}) \right|}{\sqrt{\langle|X_{i}(f_{1})X_{j}(f_{2})|^{2}\rangle\langle|X_{k}(f_{1}+f_{2})|^{2}\rangle}}.$$

For a purely real instantaneous mixing transfer function the imaginary part of $B$ is identically zero, so any non-zero $b_{anti}$ cannot be explained by leakage. In our analyses this is the reviewer-reserve leakage-immune metric; the primary measure in the main text is GCMI [9] for its established use in non-parametric M/EEG cross-frequency analysis and its numerical stability at small sample sizes.

#### S1.5 Gaussian-copula mutual information

For two continuous variables $X,Y$ with empirical ranks $r_{X},r_{Y}$ across $n$ samples, the copula transform is

$$\tilde{X}=\Phi^{-1}\left( \frac{r_{X}}{n+1} \right), \tilde{Y}=\Phi^{-1}\left( \frac{r_{Y}}{n+1} \right),$$

where $\Phi^{-1}$ is the inverse standard Gaussian CDF. The Gaussian-copula mutual information is the Shannon mutual information of a bivariate Gaussian with the empirical copula correlation,

$$I_{GCMI}(X;Y)=-1/2\ln\det corr(\tilde{X},\tilde{Y}).$$

This is exact for a multivariate-Gaussian copula and a lower bound for general non-Gaussian joints. In the phase-amplitude coupling setting $X$ is the pair $(\cos\phi_{\text{slow}},\sin\phi_{\text{slow}})$, which embeds the circular slow-band phase into $\mathbb{R}^{2}$, and $Y$ is the amplitude envelope $a_{\text{fast}}$ of the fast band. The $I_{GCMI}$ of this joint is our primary phase-amplitude coupling metric.

#### S1.6 What the theory predicts, and what it does not

The non-abelian extension makes three empirical predictions tested here. First, loci exist where the three-point function with the rest of the brain is amplified relative to the average region, and we call these coupling generators. Second, the antisymmetric part of the cross-bispectrum and the GCMI-based phase-amplitude coupling estimator are both appropriate observables for detecting such loci. Third, in low-arousal states where competing cortico-cortical activity is reduced, the generator signature should be amplified relative to the awake state, and this amplification is what we can detect in practice. The theory does not make predictions about the anatomical identity of the generators. That question is empirical, and the answer in human sleep EEG is that the generators are in the medial temporal lobe and in the basal ganglia, not in the thalamus.

### S2. A reviewer reserve: three phase-amplitude coupling metrics

We pre-computed, and retain in frozen form, per-pair values under three distinct phase-amplitude coupling metrics in anticipation of reviewer objections to a single-metric framing. The three are: the magnitude bicoherence $b$, the leakage-immune antisymmetric bicoherence $b_{anti}$ of Chella et al. [16], and the Gaussian-copula mutual information $I_{GCMI}$ of Ince et al. [9] that serves as the primary metric in the main text. All three were computed on the same Welch-segmented cross-bispectrum on the same data. The GCMI analysis identified the eight MTL and basal-ganglia parcels reported in §4.1 of the main text; the $b$ and $b_{anti}$ analyses independently identified overlapping non-thalamic parcels, although with smaller surviving counts because of their different null distributions. The qualitative picture — that the surviving set is non-thalamic and anatomically concentrated in MTL and BG — is stable across the three metrics. The raw per-parcel values are retained as a frozen audit trail on request.

### S3. Per-parcel generator tables

#### Table S1. REM theta-beta GCMI per-parcel elevation (surviving FDR $q\leq0.05$)

| Rank | Parcel | Group | Ratio | CI lower | CI upper | $q$ |
| --- | --- | --- | --- | --- | --- | --- |
| 1 | HIP-head-l-lh | HIP | 2.428 | 1.614 | 3.451 | 0.0054 |
| 2 | lAMY-lh | AMY | 2.174 | 1.578 | 2.909 | 0.0054 |
| 3 | lAMY-rh | AMY | 1.972 | 1.409 | 2.641 | 0.0486 |
| 4 | HIP-body-lh | HIP | 1.926 | 1.351 | 2.661 | 0.0140 |
| 5 | mAMY-lh | AMY | 1.644 | 1.268 | 2.111 | 0.0140 |
| 6 | HIP-head-m2-lh | HIP | 1.528 | 1.217 | 1.926 | 0.0140 |
| 7 | aGP-rh | GP | 1.492 | 1.173 | 1.939 | 0.0486 |
| 8 | PUT-VP-lh | PUT | 1.434 | 1.178 | 1.743 | 0.0261 |

Ratios are the mean AnPhy REM GCMI divided by the mean awake rest-wake GCMI for pairs touching the named parcel. Bootstrap 95% confidence intervals (10 000 resamples, seed 0) exclude unity for every entry. The $q$ values are from BH-FDR within the REM $\times$ theta-beta family of 54 subcortical parcels; sign-flip permutation (10 000 iterations, seed 0) provides the underlying $p$ values.

### S4. Split-half null validation

#### Table S2. Null distribution of surviving-parcel count per CFC pair

| CFC pair | mean | median | 95th pctl | 99th pctl | max | $P(n\geq8)$ | observed |
| --- | --- | --- | --- | --- | --- | --- | --- |
| theta-gamma | 0.78 | 0 | 5.1 | 8.0 | 8 | 0.030 | 0 |
| alpha-gamma | 0.61 | 0 | 5.0 | 10.0 | 12 | 0.020 | 0 |
| theta-beta | 1.04 | 0 | 5.0 | 11.0 | 14 | 0.020 | 8 |
| alpha-beta | 0.96 | 0 | 6.1 | 14.0 | 16 | 0.050 | 0 |

The null was constructed by drawing 100 pseudo-sleep cohorts of size 29 from the WAND+Cam-CAN awake pool and running the identical per-parcel sign-flip permutation analysis on each. $P(n\geq8)$ is the fraction of null iterations with 8 or more FDR-surviving parcels. The observed REM theta-beta value of 8 lies at the 98.0th percentile of the null; we read this as robust but not extreme, and we refrain from numerical claims about the exact number of surviving parcels.

### S5. Spectral Granger causality rollup

The spectral Granger analysis was run on the 8 generator parcels of §4.1, 16 left anterior temporal cortical targets, and 14 autonomic-or-derived channels (10 raw plus 4 ECG-derived respiratory surrogates), in three bands (infraslow, theta, beta), with band-adapted epoch lengths (60 s with 50% overlap for infraslow, 20 s non-overlapping for theta and beta), and per-pair paired t-tests with BH-FDR correction within each (stage, band, direction-pair) family.

#### Table S3. Granger rollup

| Stage | Band | Direction pair | Survivors | Dominant group (sign) |
| --- | --- | --- | --- | --- |
| N1 | infraslow | auto $\to$ gen | 24 / 64 | cardiac (+); EDR-A (−) |
| N1 | theta | auto $\to$ gen | 23 / 64 | cardiac (+) |
| N1 | beta | auto $\to$ gen | 24 / 64 | cardiac (+) |
| N1 | beta | gen $\to$ cortex | 3 / 128 | HIP $\to$ left anterior insula |
| N2 | infraslow | auto $\to$ gen | 25 / 64 | cardiac (+); RSA (−) |
| N2 | infraslow | gen $\to$ cortex | 20 / 128 | HIP / AMY $\to$ limbic-A temporal pole |
| N2 | theta | auto $\to$ gen | 26 / 64 | cardiac (+), $t=+6.0$ best |
| N2 | beta | auto $\to$ gen | 25 / 64 | cardiac (+), $t=+5.2$ best |
| REM | infraslow | auto $\to$ gen | 32 / 64 | EDR-A / RSA (−) dominant |
| REM | theta | auto $\to$ gen | 24 / 64 | cardiac (+), $t=+5.4$ |
| REM | theta | gen $\to$ cortex | 20 / 128 | cortex $\to$ hub (all 20 negative) |
| REM | beta | auto $\to$ gen | 25 / 64 | cardiac (+), $t=+5.6$ best |
| REM | beta | gen $\to$ cortex | 24 / 128 | HIP / mAMY / PUT-VP $\to$ insula, DMN, TempPole |

Survivors are the pair-level BH-FDR survivors within each (stage, band, direction-pair) family. Signs on groups indicate the sign of $(forward GC-reverse GC)$ mean across FDR-surviving pairs: $+$ means the autonomic side or the generator side leads.

#### S5.1 EDR-A and RSA in the infraslow band

Splitting the “respiratory-derived” autonomic sub-group into three methods (EDR-A, EDR-BW, RSA) reveals a consistent pattern at infraslow frequencies. In all three sleep stages EDR-A and RSA show the hub $\to$ envelope direction with comparable magnitude and $t$ value. EDR-BW shows the opposite direction, aligned with the raw ECG, because it shares low-frequency content with the cardiac waveform itself and is therefore not a clean mechanical control. EDR-A is a purely mechanical surrogate, derived from R-peak amplitude modulation during thoracic-axis rotation with breathing, and does not use the R-R intervals at all; its agreement with RSA argues against the simplest null hypothesis that the hub $\to$ envelope finding is reducible to the hub controlling heart-rate variability. Reading the rollup this way, the data are consistent with the hub leading the slow respiratory envelope in the infraslow band in all three sleep stages we examined.

### S6. Quality control and exclusions

Exclusions were applied on pre-specified quality criteria before the analyses below were run. Sleep subjects with fewer than 10 qualifying N1 epochs were excluded from the N1 analysis (7 exclusions, giving $n_{N1}=22$). Sleep subjects with fewer than 20 qualifying N2 or REM epochs were excluded from the respective stage analysis (0 exclusions, $n_{N2}=29$, $n_{REM}=29$). EEG channels with kurtosis $>10$ or variance more than 5 MAD from the median were flagged as bad and interpolated; sleep subjects with more than 10 bad channels were excluded from source reconstruction (2 exclusions of that kind). Sleep subjects whose ECG had fewer than 6 detectable R-peaks per 30-second epoch were excluded from the EDR analysis (0 exclusions).

### S7. Alternative-parcellation robustness

The results in §4 of the main text are conditioned on a specific choice of cortical parcellation (the 800-region Schaefer 17-network atlas) and a specific choice of subcortical parcellation (the 54-region Tian S4 atlas). A robust interpretation of the generator findings requires that the MTL / basal-ganglia concentration is not an artefact of either choice. We flag three natural alternative-parcellation checks as follow-up work:

1. **Cortical granularity.** The Schaefer atlas is released at 100, 200, 300, 400, 500, 600, 700, 800, 900 and 1000 parcels. Re-running the full §4 pipeline at 400, 600 and 1000 parcels should leave the subcortical generator identification qualitatively unchanged, because the subcortical test does not depend on the cortical parcel set. The cortical-target identification (the network enrichment in limbic-A temporal pole and default-mode-B temporal cortex reported in §4.2) is expected to change quantitatively with parcel size: coarser variants merge the functional subdivisions and finer variants fragment them. A stable pattern at 400 and 1000 would support the 800-parcel choice in the main text.
2. **Subcortical granularity.** The Tian subcortex atlas is released at four scales (S1, S2, S3, S4), corresponding to 16, 32, 50 and 54 parcels. Re-running the generator identification at S2 and S3 should collapse adjacent hippocampal and amygdala subdivisions without changing the anatomical class of the surviving parcels. A stable qualitative picture across S2, S3 and S4 would support the S4 choice.
3. **Independent atlases.** The Brainnetome atlas is a plausible independent alternative that includes both cortical and subcortical regions at a single granularity. Applying the §4 pipeline to the Brainnetome parcellation, on the same data, would provide a strong cross-atlas replication of the MTL and basal-ganglia generator identification.

These checks are not performed in the present paper. They are a natural next step and we flag them explicitly so that independent groups can undertake them.

### S8. VAia as a nearby non-Re thalamic negative control for the Morel Re analysis of §4.5

Section 4.5 of the main text extracts a Morel-defined nucleus reuniens (Re) time series at 1 mm source resolution and reports a directed PFC-Re-MTL signature that matches the bands and direction predicted by the $G_{e}$ commitment of §2 and by the rodent lesion result of Kafetzopoulos et al. [52]. The question this supplementary section addresses is whether that signature can be explained away as beamformer point-spread from neighbouring ventral-anterior thalamic tissue, rather than as a genuine reuniens contribution. The answer is no, and the test that gives that answer is a side-by-side extraction of the bilateral Tian S4 ventral-anterior inferior-anterior thalamic parcel (THA-VAia) as a negative control, carried through identically the same spectral-Granger pipeline as Morel Re in §4.5.

**Why VAia is the correct negative control.** The Morel Re mask [53, 54] is centred at the ventral midline of the thalamus (approximate MNI $(-1.9,-8.8,-1.9)$ on the left and $(+3.3,-8.5,-1.6)$ on the right) and occupies about 16 mm${}^{3}$ per hemisphere. The Tian S4 THA-VAia parcel sits about 5.3 mm lateral, dorsal and slightly posterior, overlaps only $\sim1\%$ of Morel Re voxels by intersection, but is close enough that at the $\sim5$ mm subcortical source grid used in the main pipeline a beamformer point-spread function centred on Re would reach well into VAia. A genuinely Re-localised signal in the human EEG should therefore partially leak into a VAia extraction through the same LCMV filter, with a reduced amplitude because of the off-centre point-spread weight. In the opposite direction, a signal whose true anatomical origin is inside the VAia mask but outside Re — for instance a ventral-anterior thalamic contribution unrelated to the reuniens-PFC-hippocampus circuit — would show up in the VAia extraction and not (or only partially) in the Re extraction, because Re sampling is restricted to the 29 Morel voxels and does not reach the VAia centroid. The negative-control logic is therefore symmetric: the shared cells show beamformer-shared tissue, the Re-exclusive cells localise signal to tissue inside Re and outside VAia, and the VAia-exclusive cells (of which there are none in the present results) would localise signal to tissue inside VAia and outside Re.

**Pipeline parity.** VAia was carried through Phase A and Phase B of the same run_granger_morel_pfc.py pipeline reported in §4.5. The only difference is the seed: Morel Re uses the 29 densified 1 mm voxels, averaged per hemisphere, and VAia uses the Tian S4 parcel average per hemisphere. Target sets (8 MTL / BG generators, 68 prefrontal parcels, 16 left anterior-temporal parcels, and the two raw cardiac channels), bands (infraslow 0.05-0.5 Hz, theta 4-8 Hz, beta 13-30 Hz), epoch lengths (60 s infraslow, 20 s theta and beta), the Dhamala non-parametric state-space Granger solver, and the per-pair BH-FDR correction within each (stage, band, direction-pair) family, are identical.

**Cells shared by Re and VAia: the expected point-spread floor.** In N2, across all three bands tested, VAia reproduces roughly two-thirds of the Re-to-PFC directed couplings with near-identical effect sizes at the top pairs. In N2 infraslow the top Re-to-OFC pair was Morel_RE-rh $\to$ LH_LimbicB_OFC_11-lh (forward-minus-reverse $+0.0481$, $t=+5.62$, $q=0.0007$); the matching VAia pair was THA-VAia-rh $\to$ LH_LimbicB_OFC_11-lh (forward-minus-reverse $+0.0484$, $t=+6.24$, $q=0.0001$). The agreement at the top-pair level is consistent with the expected partial-volume mixing between the two extractions and should not be read as a VAia-specific effect. Similarly, N2 theta (13 Re-to-PFC pairs, 6 VAia-to-PFC pairs) and N2 beta (23 Re-to-PFC pairs, 13 VAia-to-PFC pairs) both show Re dominating VAia in surviving pair count while the top pairs are nearly identical between the two extractions. We read the N2 prefrontal cells as jointly Re-driven with an unavoidable VAia contamination from point-spread, and we do not claim they are purely Re-specific.

**Re-exclusive cells: evidence against point-spread.** Three cells in §4.5 survive the negative-control test in the strong sense, with zero VAia counterparts:

1. **N2 infraslow Re** $\to$ **temporal pole**: 5 Morel Re pairs surviving FDR correction (top: Morel_RE-lh $\to$ LH_LimbicA_TempPole_6-lh, $t=+4.99$, $q=0.0009$), and 0 VAia pairs in the same cell. This cell provides the strongest individual evidence for a Re-specific contribution in the analysis.
2. **N1 beta Re** $\to$ **prefrontal**: 2 Morel Re pairs surviving FDR correction (Morel_RE-rh $\to$ RH_LimbicB_OFC_9-rh, Morel_RE-rh $\to$ RH_DefaultA_PFCm_3-rh), and 0 VAia pairs.
3. **N1 beta Re** $\to$ **temporal**: 2 Morel Re pairs surviving FDR correction, and 0 VAia pairs.

The argument against point-spread rests on the symmetry of beamformer leakage. If the underlying signal lived in the shared Re+VAia tissue rather than specifically in Morel Re, the leakage would have split between the two extractions, and both would have shown surviving pairs (the specific split depending on point-spread weights, but never all-to-none). Conversely, the observed all-to-none pattern is consistent with signal originating in tissue inside Morel Re and outside the VAia mask — i.e., inside the reuniens. The three cells therefore pass the negative-control test in its strong form, and we treat them as providing the strongest evidence for Re-specificity in the analysis of §4.5.

**REM beta: the one cell we do not claim as Re-specific.** In REM beta, Morel Re and THA-VAia are nearly indistinguishable by the same test: 15 Re pairs surviving FDR (top: Morel_RE-lh $\to$ LH_DefaultB_Temp_2-lh, $t=+3.19$, $q=0.0160$), 14 VAia pairs surviving FDR in the same direction, matching at the top-pair level in both sign and effect size. The negative-control test cannot distinguish Re from VAia here. The honest reading is that REM beta carries a broad ventral-midline-thalamic to anterior-temporal-cortex signal that cannot be dissected at the source resolution available from 71-channel scalp EEG. Main text §4.5 reports this as a limitation rather than as a Re-specific cell, and the consistency with the framework-committed prediction rests on the three unambiguous Re-specific cells above, not on REM beta.

**VAia-exclusive cells: none.** We also looked for cells where VAia surviving pair count exceeds Re and the top VAia pairs are absent from the Re extraction. In the scan across all three stages, three bands and twenty directed blocks, no such cell was identified. The only VAia-leaning cell in the full rollup is REM theta temporal (5 VAia pairs vs 4 Re pairs), where the two extractions agree to within a single pair and the top pairs are shared. We do not interpret this as VAia-exclusive.

**Summary.** The Morel Re extraction of §4.5 passes a strict negative-control test against a neighbouring thalamic parcel that shares about 5 mm of spatial separation and would inherit any non-trivial Re beamformer point-spread. The three cells we identify as possibly Re-specific (N2 infraslow Re-to-temporal-pole and the two N1 beta Re-to-prefrontal / Re-to-temporal cells) show zero VAia counterparts, a pattern that argues against point-spread as the sole explanation and is consistent with tissue-localised Re activity. The shared N2 prefrontal cells remain in the main text as jointly Re- and VAia-reflecting signals that are consistent with the broader ventral-midline-thalamic-to-prefrontal routing suggested by the framework but cannot be attributed exclusively to Re. REM beta is reported as an equivocal cell.

### Supplementary references

The supplementary material cites the same reference set as the main text; see main-text References [1]–[102].
